# MIC*: A Framework for Interpretable Analysis of Ordinal Viability Data

**DOI:** 10.1101/2025.10.13.682067

**Authors:** Carson L. Stacy, Sonali Lenaduwe, Tara N. Stuecker, Jeffrey A. Lewis

## Abstract

Microbial survival assays frequently use ordered categorical (ordinal) scores, such as semi-quantitative viability scores across drug concentrations. While these ordinal data provide rich information about dose-response dynamics, they require appropriate statistical approaches for proper analysis. Proportional-odds (PO) ordinal regression is specifically designed for such data, modeling the ordered nature of scores while accommodating continuous variables such as concentration. However, despite its advantages, PO regression remains underutilized in microbiology because its cumulative log-odds outputs are abstract and biologically unintuitive. Consequently, researchers often resort to flawed alternatives: collapsing scores into binary outcomes (growth/no growth), treating scores as continuous values for t-tests or ANOVA, or applying nonparametric tests that ignore dose-response structure. Each approach sacrifices power or validity, risking unreliable conclusions. To allow researchers to better leverage the power of ordinal datasets, we introduce MIC*, a summary measure that translates PO regression results into biologically interpretable units. MIC* is defined as the treatment concentration where the predicted probability of “no viability” equals 0.5, and is thus conceptually related to the minimum inhibitory dose (MIC) widely used in antimicrobial research. MIC* retains the rigor of ordinal regression while providing more biologically intuitive effect size measures. The MIC* framework enables formal comparisons as absolute differences (ΔMIC*) or relative fold-changes (Δlog_2_MIC*), allowing for robust statistical comparisons between samples. Monte Carlo simulations demonstrate that MIC* yields robust estimates with superior power compared to conventional statistical methods, while case studies demonstrate practical utility. To ensure wide adoption, we provide a suite of open-source tools: a BLInded Scoring System (BLISS) for generating ordinal viability scores, and web-based (MICalculator) and R package (ordinalMIC) alternatives for performing complete MIC* analyses, thus reducing barriers to appropriate analysis of ordinal phenotypes in microbiology.

**Importance:** Microbiologists routinely use visual scoring systems to assess cell viability for antimicrobial and stress resistance studies. However, current analysis approaches tend to be problematic, with researchers seeking simple approaches that either discard valuable information or violate important statistical assumptions. While rigorous and appropriate statistical approaches exist, they are rarely used in microbiology because they produce outputs that are non-intuitive and require specialized expertise to implement. Our MIC* framework overcomes these longstanding barriers by translating abstract outputs into an intuitive, concentration-based metric akin to the minimum inhibitory concentration (MIC) well Known to microbiologists. MIC* allows for biologically interpretable comparisons on both absolute (ΔMIC*) and relative (Δlog_2_MIC*) scales, allowing for facile between-sample comparisons. MIC* outperforms conventional statistical approaches, and we offer user-friendly software tools to enable broad adoption by the community.

## Introduction

Quantifying cellular survival is a key aspect of modern microbiology research, with applications ranging from clinical antimicrobial susceptibility to food safety testing and drug discovery. A key parameter of interest is the minimum inhibitory concentration (MIC), traditionally defined as the lowest concentration of a compound that prevents visible growth, or in the context of survival assays, the concentration that reduces viability to a specified threshold. Gold-standard assays for MIC determination, such as broth or agar dilution, provide valuable insights into concentration-dependent survival (1, 2). However, these methods are less amenable to high-throughput quantification compared to growth assays. For example, MIC determination via broth microdilution or disc diffusion techniques are widely used, but are relatively low-throughput and labor-intensive, typically yielding a single data point per sample. This limitation is particularly acute when the goal extends beyond simply classifying a strain as resistant or susceptible, but is instead to precisely quantify biologically relevant phenotypic differences. High-throughput plate-based “spot assays” that score viability offer a promising solution, allowing for the rapid screening of many isolates across a full dose-response series (3, 4), generating rich datasets with viability scores at multiple concentrations for each sample. However, the potential insights obtained from these rich datasets are rarely realized, as current analyses generally collapse the dozens of score spots into a single MIC endpoint. Collapsing the data discards valuable information and reduces statistical power to detect meaningful biological variation.

While sophisticated methods for analyzing high-throughput assays of microbial survival have been developed, their implementation presents practical and statistical challenges (3–12). In terms of viability, direct determination of colony-forming units (CFUs) for every spot across multiple doses is an existing alternative (13), but prohibitively resource-intensive at scale. Fully quantitative approaches, such as those based on automated imaging and spot pixel intensity analysis, can be powerful but require considerable computational expertise, model training, and are sensitive to imaging artifacts (4–7, 14). Microplate-based growth rate determination provides a quantitative alternative, but can be hampered by many sources of technical variation (e.g., plate type, shaking speed, aeration, edge effects), and requires specialized equipment for high-throughput applications (15). Thus, classical MIC determination methods lack informational richness suitable for high-throughput analyses, while more sophisticated alternatives remain difficult to implement accurately, limiting broader adoption.

Facile measurement of survival with greater phenotypic detail is already available via semi-quantitative scoring of viability (16, 17). By incorporating researcher expertise to assign scores to represent the degree of viability at each concentration, rich information about cell survival is obtained while controlling for experimental variation that is difficult to automate. This scoring approach is amenable to low, medium, and high-throughput experimentation (Figure 1), and provides substantially more information than simple binary (i.e., growth/no growth) classifications (18). However, the assigned scores are ordinal in nature, meaning they represent natural ordered categories (e.g., 0 < 1 < 2 < 3), but the “distance” between scores is not necessarily equal. Thus, semi-quantitative scores require analyses that are appropriate for ordinal data, but many biologists lack the statistical tools for analyzing ordinal phenotype data across experimental groups and concentrations. As a result, researchers are often forced to choose between flawed analytical shortcuts (Table 1) or sacrifice statistical power to detect observed differences. For example, treating ordinal scores as equally spaced numbers for use in a *t*-test or ANOVA relies on a biologically suspect assumption (equal spacing across scores) that can inflate false positive rates (19, 20). Conversely, collapsing spot scores into a binary (growth/no growth) outcome discards valuable phenotypic information, thereby reducing statistical power to detect important biological differences (21).

**Figure 1.**
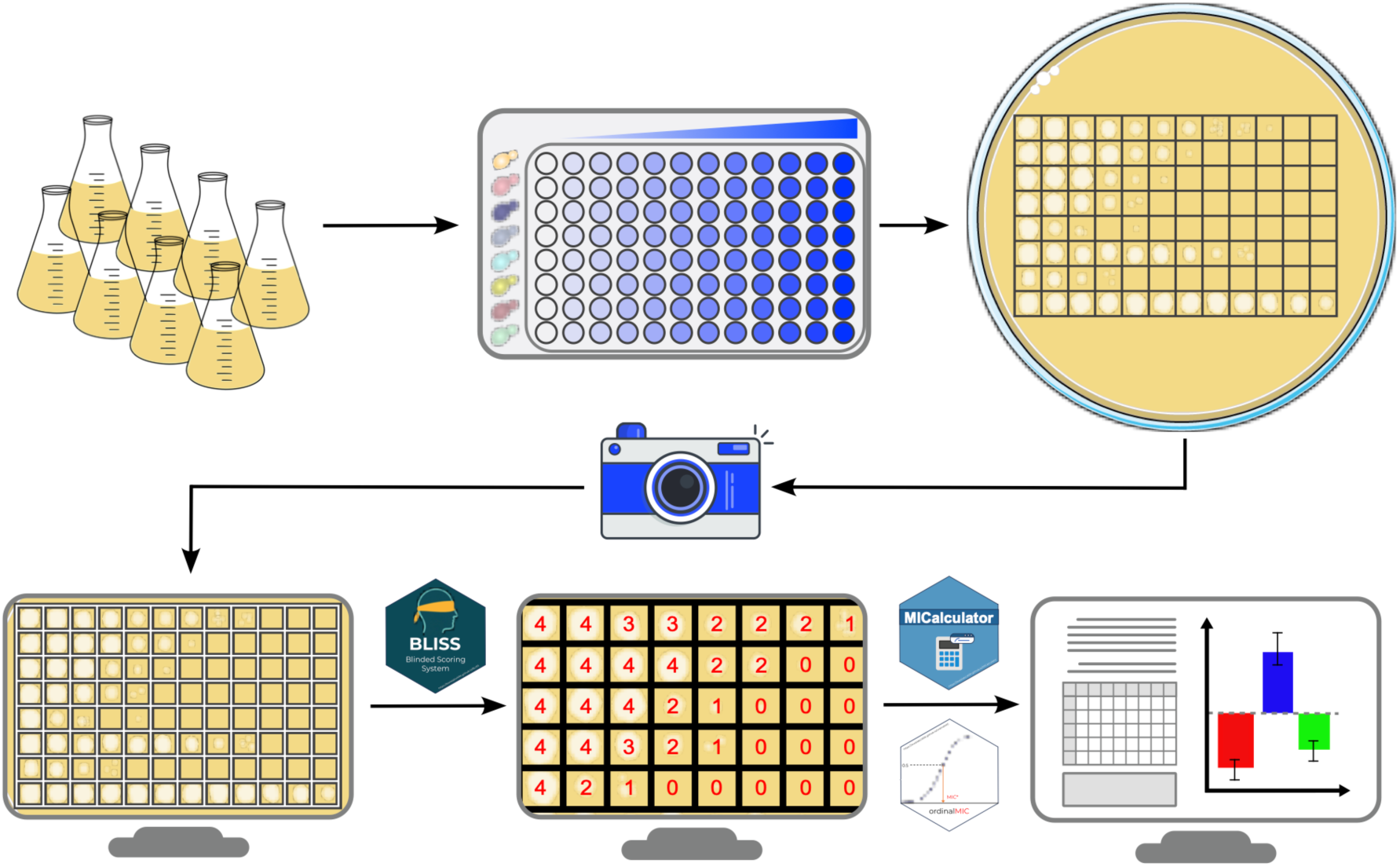
Workflow for semi-quantitative viability experiments, blinded scoring (BLISS), and MIC* analysis. Cells are assayed in 96-well plate format across an 11-dose concentration series of an experimental treatment (e.g., stressor or drug), plus untreated controls. After imaging, the BLInded Scoring System (BLISS) software segments plate images, randomizes spots for blinded scoring, and exports MIC* analysis-ready files. MICalculator or ordinalMIC software then fit PO models to estimate MIC* (the treatment concentration where the probability of no observed viability is ≥50%) and perform statistical comparisons between experimental groups.

**Table 1:** Comparison of common analytical approaches for ordinal dose-response data.

| Analytical Approach | Description | Key Assumption(s) | Primary Limitation(s) | Key References |
| --- | --- | --- | --- | --- |
| *t*-test of Observed MIC | Dose-response of each replicate is summarized by MIC value. A *t*-test compares means of MICs between groups. | MIC values are continuous, normally distributed, and vary between replicates. | Discards dose-response information (major loss of power), violates statistical assumptions as MICs are interval-based, not continuous. | (17, 20, 22, 23) |
| Dichotomization ($\chi^2$, Fisher's exact) | Growth at each concentration is classified as binary (e.g., +/-). Proportions at each concentration are compared between groups. | Intermediate growth phenotypes are biologically irrelevant; only distinction between two chosen categories matters. | Massive loss of statistical power by discarding dose-response information. Results depend on the chosen cut-off. | (24) |
| *t*-test / ANOVA on Scores | Ordinal scores (e.g., 0-4) are treated as continuous values. The mean score (or sum of scores) is compared between groups. | Scores are biologically equidistant (e.g., two scores of 1 = one score 2). | Imposes a false interval scale on ordinal data, violates core statistical assumptions and potentially inflates the rate of false positives. Treats all concentrations equally. | (25–27) |
| Nonparametric tests (e.g., Wilcoxon) | Ranks of ordinal scores are compared between groups without assuming a data distribution. | Only rank order of outcomes is meaningful, values of concentration are irrelevant | Lower statistical power than regression. Cannot incorporate continuous predictors; ignores dose-response. Equivalent to PO model with only categorical variables. | (28) |
| Proportional-Odds Regression model | Models the cumulative probability of ordinal scores as a function of both categorical (e.g., genotype) and continuous (e.g., concentration) predictors. | The effect of predictors is consistent across different thresholds between outcome categories. Response has a natural order. | Primary output (log-odds ratio) is non-intuitive and difficult to interpret biologically. Violation of PO assumption can lead to biased estimates. | (29, 30) |
| ΔMIC* | The absolute difference in model-derived MIC* between experimental groups (e.g. MIC*_mut - MIC*_WT) | An *additive* effect model is appropriate; the *absolute* change in concentration is the most biologically relevant measure of difference. | Provides a direct, interpretable measure of change in original units of predictor (e.g., uM). Best suited for assessing absolute potency | This Paper |
| $\Delta\log_2\text{MIC}^*$ | Log fold change in MIC between groups, unitless | A *multiplicative* effect model is appropriate; the *relative* fold change in concentration is the most biologically relevant measure of difference. | Provides a unitless, symmetric measure of relative change interpretable by MIC* change. Particularly useful for comparing effects across different compounds or skewed MIC distributions common in microbiology. | This Paper |

Fortunately, a statistically rigorous method for analyzing ordinal response data exists: proportional-odds (PO) ordinal regression (31–33). The PO model is a type of cumulative-logit regression that models relationships between an ordinal response variable and one or more predictor variables. In the context of survival assays, ordinal spot viability scores would be the response variable, and predictor variables could include strain genotype, antibiotic/drug concentration, treatment conditions, and their interactions. The PO model properly incorporates the ranked nature of the scores without assuming they are equally spaced, but wider adoption by experimental biologists has been hindered by the difficulty of application and interpretability (34). For example, the results of the PO model are expressed as *β* coefficients that represent changes in cumulative log-odds ratios, a mathematical abstraction that is biologically non-intuitive (35). Additionally, the need for specialized statistical expertise places PO ordinal regression largely out of reach for many biologists who could benefit from its use (30, 36). These challenges have been recognized, and alternative measures for reporting effects have been proposed (34, 37). However, these measures only work well for comparing groups (e.g., wild-type versus mutant), while poorly quantifying impacts of numeric variables important to biologists (e.g., dose-response concentrations).

We were thus motivated to make PO ordinal regression more accessible to biologists by translating its complex outputs to an intuitive, biologically scaled metric. Our framework uses a fitted PO ordinal regression model to derive a threshold estimate on the scale of a continuous covariate across experimental (categorical) groups. To provide a concrete application in microbiology, we illustrate this concept as a model-derived Minimum Inhibitory Concentration, or MIC*. We define MIC* as the estimated drug or stressor concentration at which the probability of observing “no viability” equals 0.5 (Figure 2A-C). In other words, MIC* estimates the median concentration where no survival would be observed in exactly half of replicate experiments. This calculation translates the abstract output of the ordinal regression model into an estimated MIC-like value that is expressed in units of the experiment (e.g., *μM* of antibiotic). Importantly, our framework also includes a robust method for estimating the variance of MIC*, allowing for direct statistical comparison and hypothesis testing between experimental groups. While we use the term MIC* for its broad recognizability, our framework is equally applicable to estimating both inhibitory and -cidal thresholds, depending on experimental design. We demonstrate the utility and power of this approach via extensive Monte Carlo simulations and empirical case studies of microbial viability applied to *Saccharomyces cerevisiae* (brewing yeast) stress responses. To ensure broad adoption, we provide open-source browser-based software for blinded viability scoring (BLISS) and MIC* analysis (MICalculator). We also provide an R package (**ordinalMIC**) implementation of the analytical methods described here.

**Figure 2.**
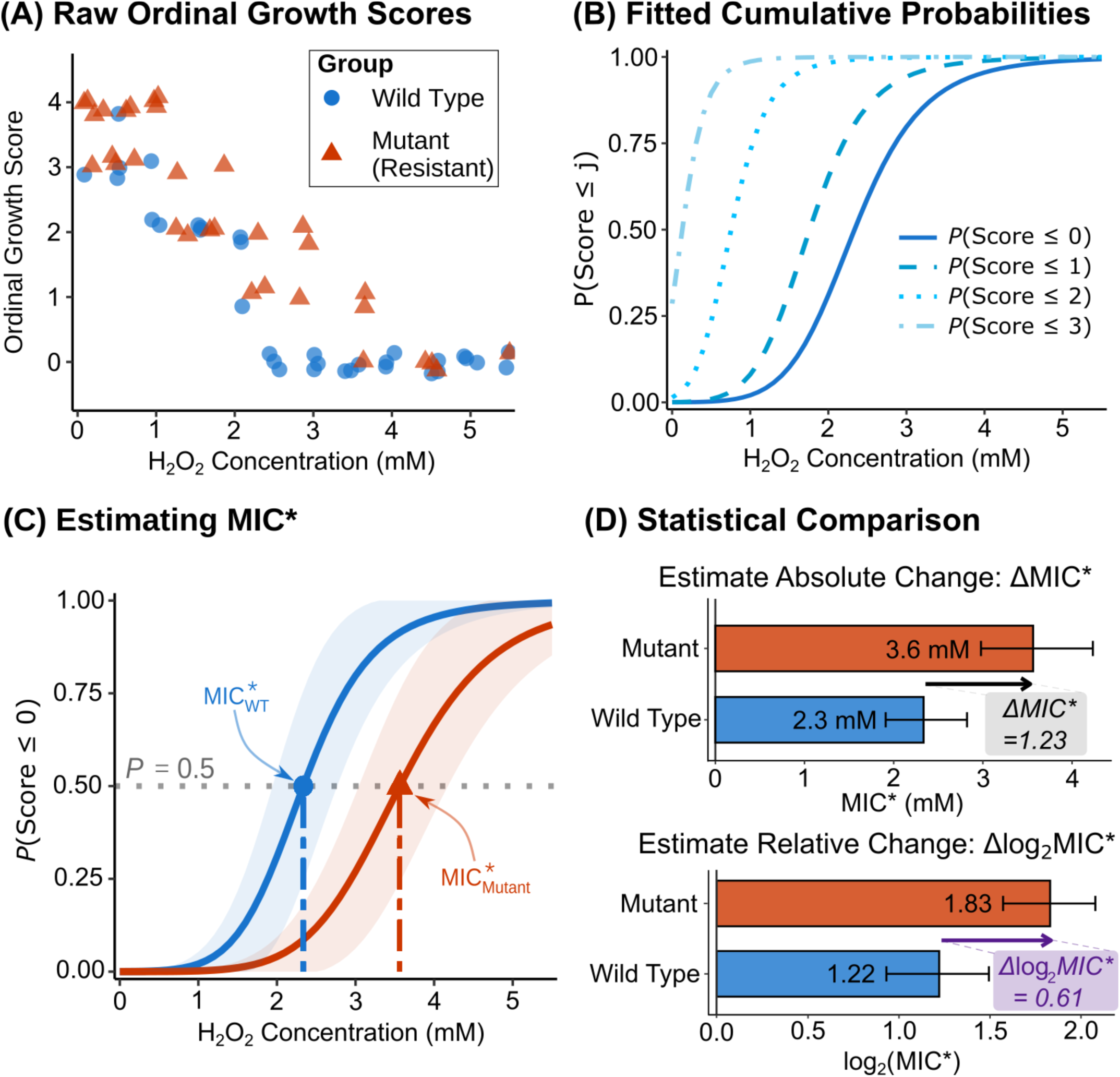
Conceptual schematic of model-derived minimum inhibitory concentration (MIC*) estimation for ordinal viability data. **(A)** Simulated raw ordinal viability scores (score 0: no viability to score 4: maximal viability) across a range of stressor concentrations for hypothetical wild-type and stress-resistant mutant strains.. **(B)** PO regression model fit yields cumulative probabilities P(Y ≤ *j*), which represent the probability of a viability score less than or equal to score *j* (0, 1, 2, 3) across stressor concentrations. **(C)** MIC* is defined as the stressor concentration at which the predicted probability of observing no viability (i.e., a score of 0) is 0.5. The MIC* estimates for the wild-type and stress-resistant mutant strains are shown, with shading representing the 95% confidence intervals (CIs). **(D)** Group comparisons for ΔMIC* (absolute difference in MIC*) and Δlog_2_MIC* (relative difference in MIC*). Error bars and shaded regions represent model-derived 95% CIs.

## Results

### The MIC* Framework

PO models are well-suited to ordinal viability data, but the resulting *β* coefficients are difficult to interpret biologically. For viability data, we propose MIC* as an easily interpretable alternative. Our approach takes ordinal viability scores, and then fits an ordinal regression model to the entire dose-response dataset (process shown in Figure 2A-B). Instead of *β* coefficients, the primary output is MIC*— formally defined as the estimated concentration at which the probability of observing “no viability” equals 0.5 (Figure 2C). By translating model coefficients into a biologically interpretable value, MIC* enables direct statistical comparisons of survival between experimental groups (Figure 2D). We facilitate group comparisons on either absolute (ΔMIC*) or relative (Δlog_2_MIC*) scales using analytical SEs and Wald tests, thereby providing biologically intuitive and statistically robust endpoints for quantifying and comparing experimental groups.

### Simulations Demonstrate High Accuracy and Power of MIC*

We first evaluated the MIC* estimator using a factorial simulation layout to mirror realistic microbial experiments in which strain- and condition-specific survival is tested over a series of treatment concentrations. Specifically, each experiment comprised a 2×2 factorial design (e.g., two genotypes × two treatments), 12 stressor (or drug) concentrations, and three biological replicates (*n=144* total observations per simulated experiment) with a range of genotype and pretreatment effects on survival (Example dataset summary in Table S1). Across 1000 Monte Carlo simulations per parameter combination (see Supplemental Methods S6), MIC* estimates from proportional odds (PO) models recovered true between-group differences at high rates with acceptable rates of false positive results via hypothesis testing on ΔMIC* or Δlog_2_MIC* scales (Figure 2D). MIC* provides a biologically scaled summary of ordinal outcomes that can be directly compared in experiments.

A “Simplified” PO model with a single interaction term for the primary interaction of interest exhibited greater power and better error control than a saturated alternative. For MIC* testing, the Simplified PO model (main effects and genotype × pretreatment interaction) reached 80% power at a ≈28% reduction in survival (𝛽_𝐺×𝑃×𝐶_ = −0.332; Figure 3A), while controlling the false positive rate (≈0.037 with a target of 𝛼 = 0.05; Figure 3B). In contrast, the Saturated model (with all 2-way and 3-way interactions) required larger effect sizes (≈40% reduction), and showed an inflated false positive rate (≈0.107) at typical sample sizes with biological triplicate (Figures 3, S1, S2). The increased power of the Simplified model comes with an expected bias-variance tradeoff: MIC* point estimates showed a modest, predictable shrinkage from the true value (mean relative bias ≈16%; Figure S3) due to the existence of the three-way interaction in the simulated datasets. Meanwhile, the Saturated model produced highly accurate MIC* estimates (mean relative bias <1%, Figure S4) at the cost of statistical power and error control (Figure 3). In practical terms, these findings suggest the more parsimonious model specification is preferable for detecting and quantifying differences between groups when analyzing data in biological triplicate with ordinary sample sizes, while the saturated model, if biologically motivated, provides unbiased estimates. Hence, all subsequent analyses reported here utilize the Simplified Model structure for calculating and testing ΔMIC*.

**Figure 3.**
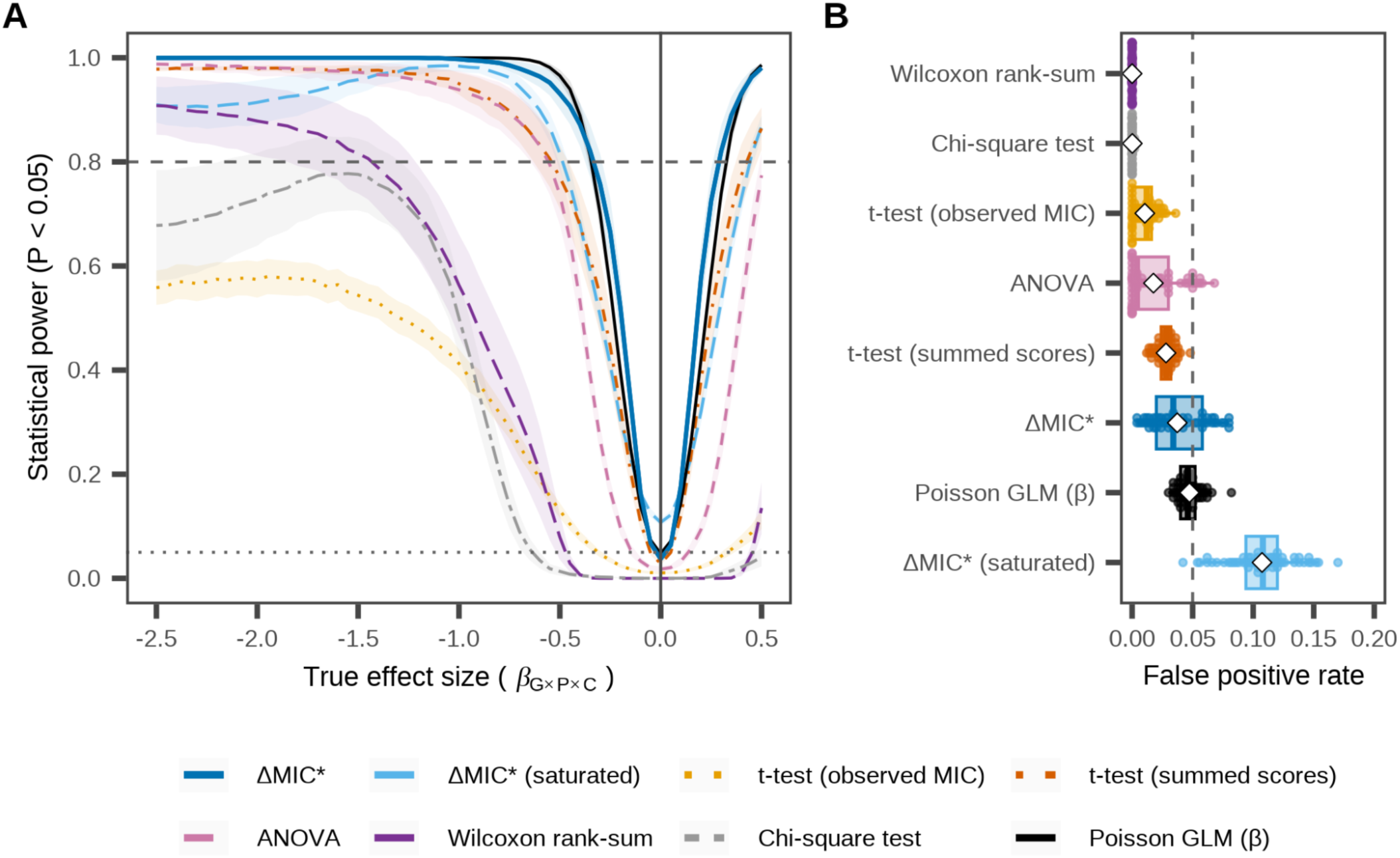
Simulations show model-derived MIC* has higher power compared to classical tests in simulations and appropriate error control. **(A)** Comparison of statistical power between ΔMIC* and traditional methods (*χ*^2^, ANOVA, t-test) across a range of simulated effect sizes. Power curves were estimated from Monte Carlo simulations (see methods), and shading represents 95% CIs across simulations. The horizontal dashed line indicates 80% power. The ΔMIC* curve shows the power of the simplified PO model. **(B)** Box plots depict observed false positive rates for each statistical method measured by simulations where no true effect existed (i.e., effect size = 0). The vertical dashed line indicates a target false positive rate (α) of 0.05.

We next compared the PO model of MIC* to ‘conventional’ statistical approaches. Compared to all other approaches tested, MIC* had the greatest power (Figure 3A). For parametric tests, a *t*-test of summed scores across biological triplicates showed moderate power to detect true differences (80% power achieved at 𝛽_𝐺×𝑃×𝐶_ = −0.54, or a ≈ 41% reduction in relative survival) while maintaining control of false positives (Figure 3B). While we ourselves have been guilty in the past of using this approach (38, 39), using a t-test to compare scores treats their labels as true numbers, which is a strong assumption. Thus, we favor nonparametric approaches appropriate for ordinal data. However, common nonparametric tests (*χ^2^*, Fisher’s exact, Wilcoxon rank-sum) required substantially larger true effect sizes to attain equivalent power for biological triplicates (≈75% reduction in survival compared to <30% for MIC*, Figure 3A). Finally, we tested Poisson models that directly analyze simulated colony count data before it was used to generate the simulated ordinal scores. The standard Poisson model represents the theoretical best-case scenario of using all colony counts, while the censored Poisson model represents the case when colony counts at the highest levels of survival are too high to count. Because these Poisson models were used to generate the simulated data set, we expected them to perform optimally. Unexpectedly, MIC* testing exhibited power closely matching the benchmark Poisson model (𝛽_𝐺×𝑃×𝐶_ = −0.34, a ≈ 29% reduction), and clearly outperformed the censored Poisson model (𝛽_𝐺×𝑃×𝐶_ ≤ −1.4 a >75% reduction). Across all testing procedures examined, MIC* was the most powerful at detecting small differences (≈30%) while controlling false positive rates, whereas the t-test, Wilcoxon, *χ^2^*/Fisher’s, and censored Poisson approaches required substantially larger effect sizes (≈40-75% reductions) to attain comparable power. MIC* retains the nonparametric robustness of the Wilcoxon test, while increasing power by leveraging the full dose-response dataset. We next sought to apply MIC* analysis to real-world viability data.

### Use Case 1: Wild Type vs Mutant Survival

For our first use case, we applied MIC* analysis to what we expect to be a common experimental design: stress survival of wild type vs mutant, either with or without a pretreatment. For this test case, we used salt-induced cross protection against H_2_O_2_ in yeast, which we know is dependent upon functional cytosolic catalase (encoded by *CTT1*) (38, 40, 41). MIC* directly translated raw semi-quantitative viability score data into easily interpretable inhibitory concentration endpoints with measured uncertainty (Figure 4). For example, in wild-type cells, NaCl pretreatment cross-protects against H_2_O_2_, shifting the estimated MIC* from 1.29 mM H_2_O_2_ (95% CI: 1.21,1.37) to 5.91 mM H_2_O_2_ (95% CI: 5.42, 6.44). The effects of pretreatment can be expressed as differences in concentrations survived (i.e., ΔMIC*), which in this case ΔMIC* = 4.62 mM, or as fold-changes in concentration survived (e.g., Δlog_2_MIC*). In this case, Δlog_2_MIC* for the effects of salt pretreatment on H_2_O_2_ cross protection is 2.20 (95% CI: 2.06, 2.33), representing a 4.58-fold increase in stressor concentration survived, indicating a statistically and biologically significant increase in MIC* (*P < 1*×*10^-10^*). In contrast to the wild-type strain, the catalase-deficient *ctt1*Δ mutant lacks NaCl-induced cross protection. This manifested as no significant effect for NaCl pretreatment: MIC*_untreated_ = 1.52 mM (95% CI: 1.44, 1.60] vs MIC*_salt_ = 1.38 mM (95% CI: 1.27, 1.51). The relative change due to salt treatment in the *ctt1Δ* mutant was quantified as a *Δ*log_2_MIC* of −0.14 (95% CI: −0.29, 0.02), which was not significantly different from zero (𝑃 = 0.09; Table 2C). We can also directly compare cross protection responses (NaCl vs Mock) of the wild-type vs *ctt1*Δ mutant strains, which in this case was a large, statistically significant difference that can be quantified in relative terms as Δlog_2_MIC*= −2.33 (95% CI: −2.51, −2.15) or in absolute difference using ΔMIC* = −4.77 mM (*P* = 2×10^-79^; Table 2). In contrast, the nonparametric Wilcoxon test on observed MICs was underpowered to detect statistical differences with biological triplicates (P = 0.1). Notably, H_2_O_2_ resistance in the absence of pretreatment does not differ between wild-type and *ctt1*Δ cells, which was captured through MIC* analysis with a Δlog_2_MIC* of 0.10 (95% CI: −0.04, 0.24; *P* = 0.17). This example represents a reanalysis of data previously summarized by the summing of viability scores (41) that recapitulates the main findings while condensing the full dose-response dataset into easily interpretable concentrations survived with confidence intervals and p-value summary statistics (Figure S6).

**Table 2A.** **MIC\*** by Strian and Pretreatment (*ctt*1Δ)

| Group |  | Estimation |  |  | Bootstrap |  |
| --- | --- | --- | --- | --- | --- | --- |
| Genotype | Pretreatment | MIC* | SE<sup>†</sup> | 95% CI | SD<sup>‡</sup> | MIC Estimates |
| WT | Mock | 1.294 | 0.025 | [1.18, 1.41] | 0.034 |  |
| ctt1 | Mock | 1.521 | 0.024 | [1.4, 1.64] | 0.053 |  |
| WT | NaCl | 5.931 | 0.042 | [5.38, 6.53] | 2.599 |  |
| ctt1 | NaCl | 1.385 | 0.027 | [1.26, 1.51] | 0.023 |  |

<sup>†</sup> SE (delta): model-based SE via delta method. **SD**: nonparametric bootstrap stratified by Genotype×Pretreatment×Concentration; 1000 successful bootstraps / 0 failed refits; B = 1000.

**Table 2B.** Pairwise contrast: ΔMIC*.

| Group |  | Estimation |  |  |  |  | Bootstrap |  |
| --- | --- | --- | --- | --- | --- | --- | --- | --- |
| Group 1 | Group 2 | ΔMIC* | SE<sup>†</sup> | 95% CI | P | FDR (BH) | SD<sup>‡</sup> | ΔMIC* Estimates |
| WT:Mock | ctt1:Mock | 0.228 | 0.074 | [0.0832, 0.372] | 0.002 | 0.003 | 0.067 |  |
| WT:Mock | WT:NaCl | 4.637 | 0.281 | [4.09, 5.19] | 6.47E-151 | 3.88E-150 | 2.604 |  |
| WT:Mock | ctt1:NaCl | 0.092 | 0.077 | [-0.0589, 0.243] | 0.232 | 0.232 | 0.046 |  |
| ctt1:Mock | WT:NaCl | 4.409 | 0.282 | [3.86, 4.96] | 3.49E-125 | 6.98E-125 | 2.597 |  |
| ctt1:Mock | ctt1:NaCl | -0.136 | 0.079 | [-0.291, 0.0196] | 0.088 | 0.105 | 0.047 |  |
| WT:NaCl | ctt1:NaCl | -4.545 | 0.284 | [-5.1, -3.99] | 5.55E-129 | 1.67E-128 | 2.596 |  |

<sup>†</sup> SE (delta): model-based SE for ΔMIC\* via delta method. **SD**: nonparametric bootstrap stratified by Genotype×Pretreatment×Concentration; 1000 successful bootstraps / 0 failed refits; B = 1000.

**Table 2C.** Pairwise contrast MIC* fold-change (Δlog_2_MIC*)

| Group 1 | Group 2 | MIC* ratio | Δlog<sub>2</sub> MIC* | SE (log<sub>2</sub>)<sup>†</sup> | 95% CI | P | FDR (BH) | SD<sup>‡</sup> | Δlog<sub>2</sub> MIC* Estimates |
| --- | --- | --- | --- | --- | --- | --- | --- | --- | --- |
| WT:Mock | ctt1:Mock | 1.176 | 0.234 | 0.076 | [0.085, 0.383] | 0.002 | 0.003 | 0.067 |  |
| WT:Mock | WT:NaCl | 4.585 | 2.197 | 0.081 | [2.04, 2.35] | 6.47E-151 | 3.88E-150 | 0.434 |  |
| WT:Mock | ctt1:NaCl | 1.071 | 0.099 | 0.083 | [-0.063, 0.261] | 0.232 | 0.232 | 0.051 |  |
| ctt1:Mock | WT:NaCl | 3.898 | 1.963 | 0.078 | [1.81, 2.12] | 3.49E-125 | 6.98E-125 | 0.427 |  |
| ctt1:Mock | ctt1:NaCl | 0.911 | -0.135 | 0.079 | [-0.29, 0.0204] | 0.088 | 0.105 | 0.044 |  |
| WT:NaCl | ctt1:NaCl | 0.234 | -2.098 | 0.085 | [-2.26, -1.93] | 5.55E-129 | 1.67E-128 | 0.424 |  |

<sup>†</sup> SE (log<sub>2</sub>): model-based SE of Δlog<sub>2</sub> MIC\* via delta method. **SD**: nonparametric bootstrap stratified by Genotype×Pretreatment×Concentration; 1000 successful bootstraps / 0 failed refits; B = 1000.

**Table 3A.**
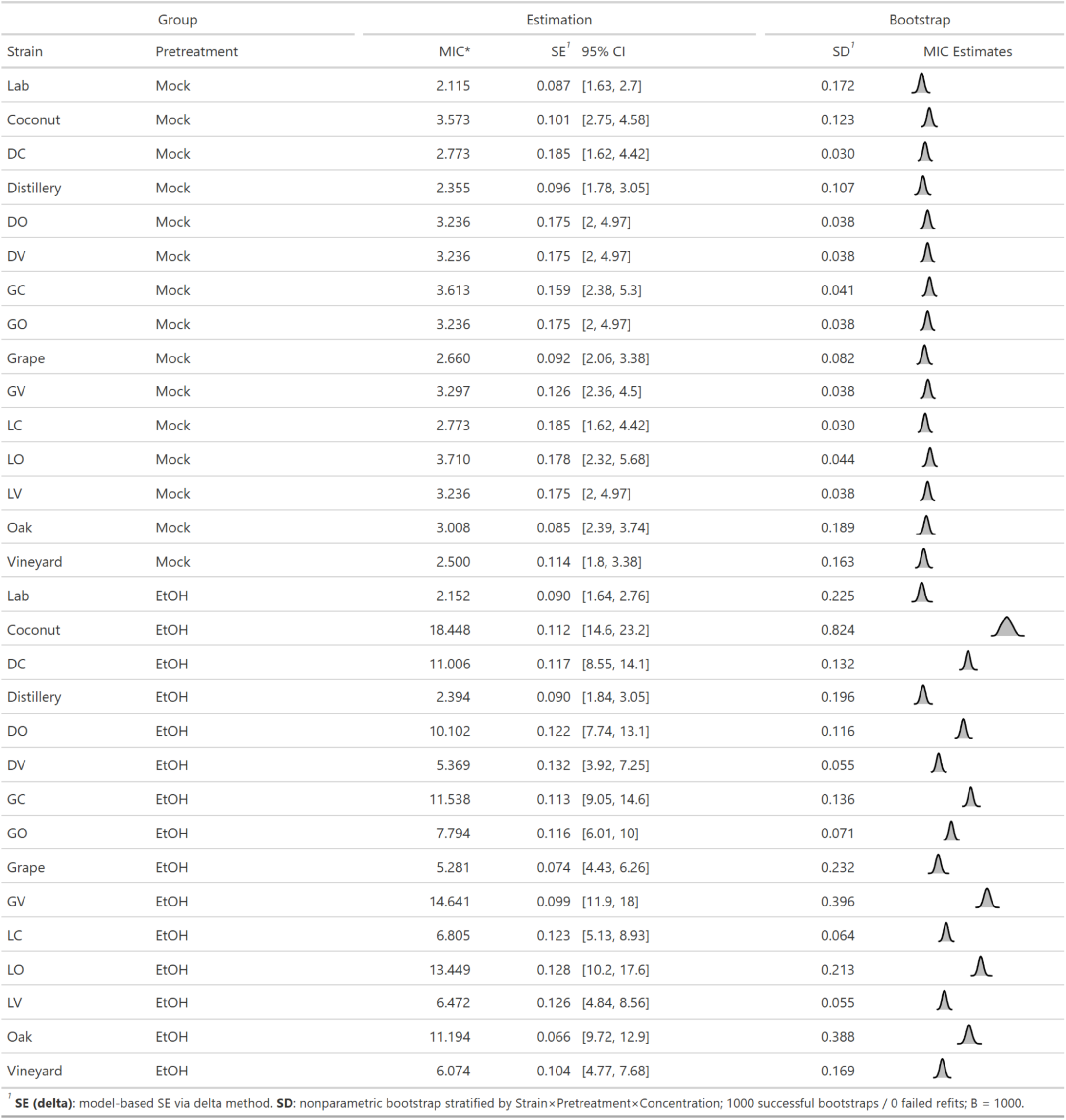
**MIC\*** of Intercross Strains

**Figure 4.**
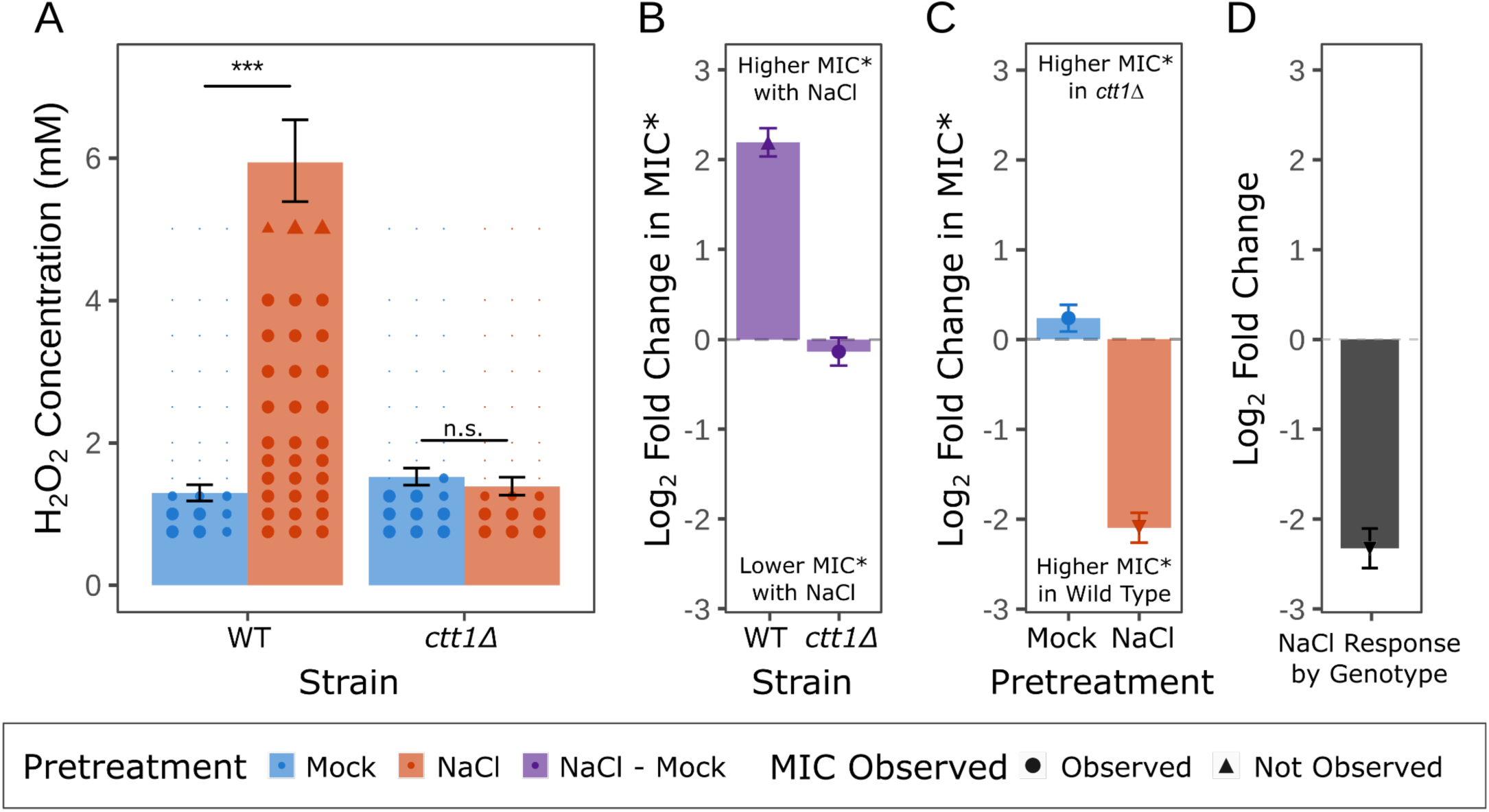
MIC* detects statistically significant differences in H_2_O_2_ survival in real experimental data with interpretable effect sizes. **(A)** The bars depict estimated MIC* for H_2_O_2_ survival for each strain, with and without mild NaCl pretreatment. Individual ordinal spot scores are shown for each experimental replicate, with triangles indicating that growth was observed at the highest tested concentration. **(B)** Effect of NaCl pretreatment on MIC* for each strain, expressed as relative fold-change (Δlog2MIC*). **(C)** Comparison of Δlog_2_MIC* between WT and *ctt1Δ* strains for each pretreatment condition. **(D)** Δlog_2_MIC* of the interaction effect showing differences in the effect of NaCl pretreatment depending on genotype. For all panels, error bars represent 95% CIs and asterisks indicate significance levels from FDR-adjusted two-sided Wald tests (* = 0.05, ** = 0.01, *** = 0.001).

### Use Case 2: Screening a Panel of Strains to Identify Differences in Stress Survival

In a second application, we used the MIC* framework to screen a panel of strains for variation in H_2_O_2_ survival with or without ethanol pre-treatment. Our motivation was to phenotype parental strains with variation in ethanol-induced cross protection, as well as progeny from advanced intercrosses derived from mating sensitive and resistant strains (Figure 5A). This test case illustrates how the improved power of MIC* can be leveraged for exploratory single-replicate (n=1) screens to more rapidly identify promising strains for more detailed replicate analysis (e.g., for quantifying doses of antibiotic survived in a panel of strains).

**Figure 5.**
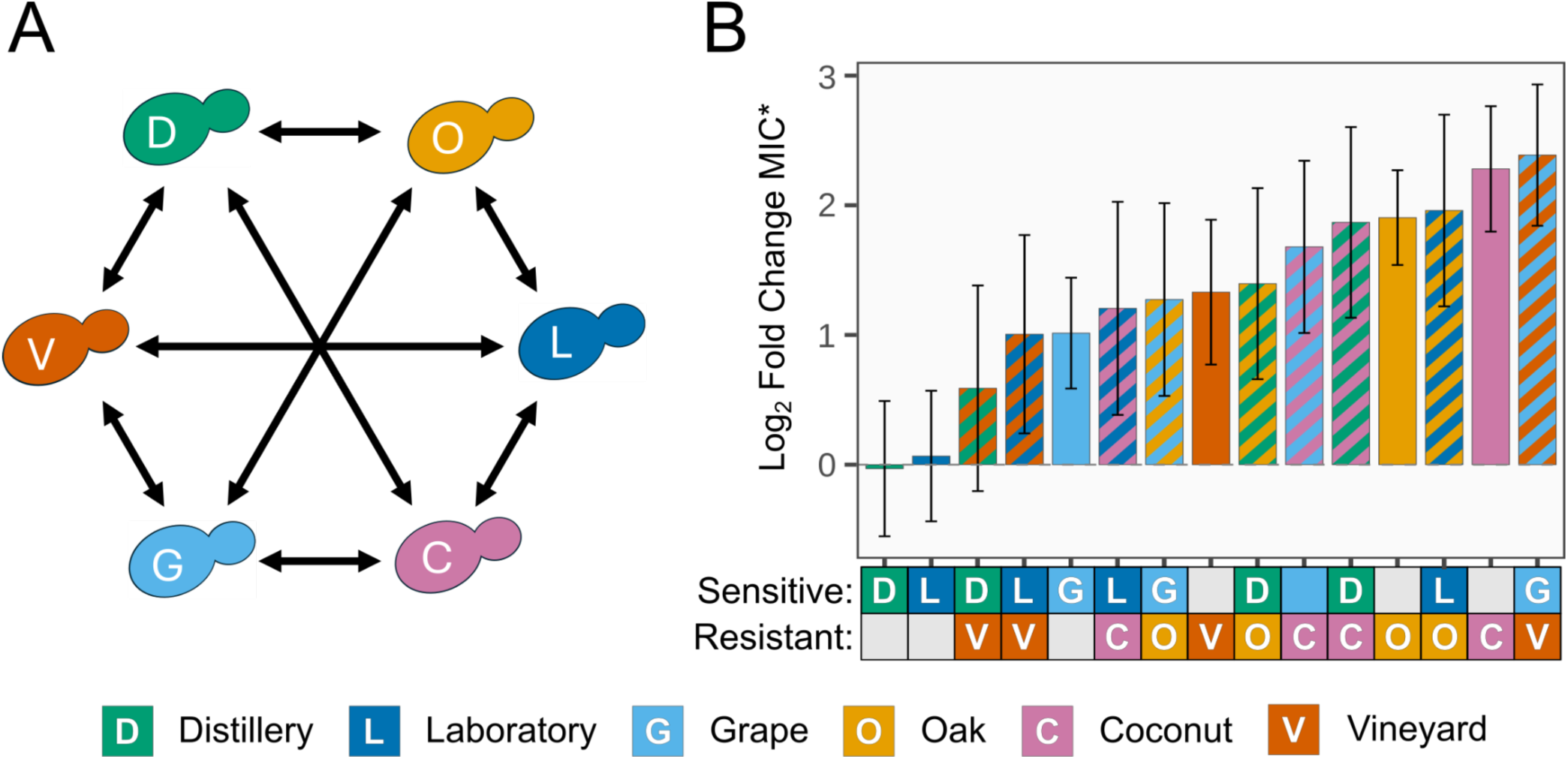
Application of MIC* to pilot viability screens quantifies phenotypic differences in yeast advanced intercross strains. **(A)** Schematic of round robin crossing design depicting all pairwise matings between H_2_O_2_ resistant and sensitive parental yeast strains. (B) Δlog2MIC* estimates showing the effect of ethanol pretreatment on H_2_O_2_ survival for parental strains and their advanced intercross progeny. Positive values indicate increased relative resistance due to ethanol pretreatment (e.g., log_2_ fold change = 1 indicates a doubling of MIC*). Solid colored bars represent parental strains, while striped bars indicate intercross progeny derived from the corresponding parental crosses. All error bars represent model-derived 95% CIs.

This pilot dataset measured H_2_O_2_ resistance in 3 resistant and 3 sensitive parental strains, as well as 9 advanced intercrosses representing every pairwise cross between sensitive and resistant strains. The dataset included both semi-quantitative ordinal scores as in Use Case 1, and also quantitative CFU counts, allowing for direct comparison of score-based MIC* and CFU count-based alternative. MIC* analysis revealed that while most intercrosses had similar H_2_O_2_ resistance in the absence of pretreatment, there was a wide distribution for ethanol-induced cross protection against H_2_O_2_. All intercrosses displayed increased cross protection relative to their sensitive parents, with the vast majority (8 of 9) estimated to have an MIC* intermediate relative to their sensitive and resistant parents. One intercross (Vineyard x Grape Strain) was estimated to display ‘transgressive inheritance,’ meaning the phenotype was more extreme than either parent, rather than being intermediate (Figure 5). Through MIC analysis of this pilot dataset, we were able to quantify between-strain differences in a preliminary screen while suggesting, with appropriate uncertainty, directions for future research.

Scores represent a simplification of some phenomena into ordered categories. In the case of cell viability, the underlying phenomenon is an intrinsically count-based (number of cells) measure of survival. As CFUs can be counted, we compared the findings of our ordinal MIC* analysis in this study to those provided with a count-based modelling approach. In this experiment, individual colony counts observed in each spot were modelled using censored Poisson regression (Figure S9). While model parameters of this Poisson regression model have a different meaning than the coefficients of the PO model, they are both formulated such that positive estimates indicate higher viability (as score or count, respectively), and lower viability when negative. Model estimates from the censored Poisson and MIC*-generating PO model exhibit highly similar estimates (parameter estimate correlation: R=0.97, Figure S10), supporting the robustness of score-based inference for pilot screens, with confirmatory replication recommended. Similar insights are derived from the example data using each model. Because this was a single replicate, the confidence intervals around MIC* are wider. Nonetheless, MIC* agreed well with CFU counts, indicating that the method is robust for preliminary screening. This result underscores the utility of ordinal modelling for characterizing phenotypic variation in microbial populations when manual counting is impractical or data are sparse.

### A Community Resource for Analysis of Ordinal Microbial Viability Data

To enable broad adoption of MIC*, we have developed several user-friendly analysis tools. First, to minimize observer bias in scoring, we developed BLISS (BLInded Scoring System), a serverless browser application that segments plate images by rows and columns into individual grid observations, randomizes presentation for blinded scoring, and records assigned ordinal scores with the option to assess scoring reproducibility with repeat scoring. Intra-scorer reliability in scoring shown here was near-perfect (weighted *κ* = 0.983), indicating near-perfect agreement (42). BLISS also exports MIC* analysis-ready data files with embedded diagnostics (Figure S11). By producing standardized analysis-ready inputs at scale from plate images through a purely browser-based workflow, BLISS separates data generation from analysis to minimize scoring bias. To empower microbiologists to perform MIC* analysis, we provide both browser-based and command-line tools for MIC* estimation and ΔMIC* testing. We developed the `ordinalMIC` R package for MIC* analysis, including fitting (via the Ordinal package), pairwise ΔMIC* testing, and visualization. By default, `ordinalMIC` reports uncertainty as a 95% CI, with standard deviation and standard error as available options. The MICalculator (LewisLabUARK.github.io/MICalculator) provides identical analysis to `ordinalMIC` in a web interface that accepts BLISS outputs (or user-provided CSV files), and returns MIC* results. Both tools perform identical analysis, facilitating MIC* estimation and comparison without coding expertise.

## Discussion

In this manuscript, we introduce MIC* as a biologically interpretable statistical framework for analyzing semi-quantitative (ordinal) viability data. MIC* takes advantage of fitted PO models designed for statistical analysis of ordinal data, while defining MIC* as the concentration at which the probability of “no viability” (i.e., a viability score = 0) is 0.5. This value acknowledges that observed MIC values are drawn from an unknown probability distribution (43). Therefore, MIC* describes the “median” minimum concentration required to eliminate all visible viability exactly half the time. Put plainly, MIC* is the dose at which, if repeated many times, about half of all runs show no viability. MIC* improves interpretability without discarding information contained in the concentration series. Unlike binarization (growth/no growth) or per-replicate endpoints (observed MIC), which discard intermediate categories and reduce power, the PO framework includes information from all scores in a single regression model. Reporting MIC*, its confidence interval, and either ΔMIC* or Δlog_2_MIC* between groups provide quantitative statements in familiar units (or unitless fold-change) that can be integrated across experiments, laboratories, and platforms through the inclusion of fixed and random effects. Absolute changes in viability (ΔMIC*) are highly interpretable as units of effect size match those of the experiment (e.g., mM drug), while Δlog_2_MIC* quantifies relative changes in MIC* that are often more relevant in studies of microbial drug resistance. Absolute comparisons provide units on the concentration scale, while when relative effects are of interest, Δlog_2_MIC* provides unitless fold-change estimates, which can be compared across treatments with different dosage ranges or units. Importantly, both ΔMIC* or Δlog_2_MIC* are derived from the same model. Further, Δlog_2_MIC* is symmetric, allowing for relative increases or decreases to be equal in magnitude and facilitating cross-study comparisons. As both approaches test the null hypothesis of whether the MIC* values for two groups are equal (MIC*_A_ = MIC*_B_), both absolute and relative results can be reported along with the corresponding measures of uncertainty. The framework generalizes to any ordered phenotype measured across a continuous variable, ranging from quantifying mitochondrial degradation to EC_50_ from biofilm scores (Supplementary Materials S13).

Our approach was theoretically motivated by the connection of PO models to rank-based non-parametric tests known to be appropriate for the analysis of ordinal data. Specifically, the PO model is a generalization of the nonparametric alternative to the *t*-test: the Wilcoxon rank-sum test (31). However, the Wilcoxon test is limited to comparing two groups, and in this simple case, the test statistic for the Wilcoxon test is a transformation of the PO model’s coefficient estimate, even when assumptions are violated (44). Importantly, the PO model, unlike the Wilcoxon test, readily accommodates additional covariates (categorical or numeric) and factorial designs, and attains much higher statistical power in small sample experimental settings involving a range of concentrations. Given this, the proportional odds assumption is important for maximizing the accuracy of MIC* estimates, but is not required for reliable statistical significance testing of group differences. In our example analyses, PO model diagnostics were consistent with no violation of the PO assumption, and confidence intervals maintained stability in simulations. MIC* estimation using PO ordinal regression incorporates information from all experimental observations, improving power while maintaining biological interpretability by quantifying effects on the concentration (or other) scale of interest. Importantly, comparison of MIC*-based testing to common alternatives suggests desirable performance for hypothesis testing. In simulations, MIC* outperformed all other standard parametric and non-parametric approaches (including the Wilcoxon rank-sum test), and approached the performance of the gold-standard Poisson model used to generate the raw counts, while clearly outperforming the censored Poisson variant. Notably, the Poisson model uses raw counts rather than ordinal scores, underscoring the power of MIC*.

Limitations exist when using MIC*, and reasonable strategies exist for mitigation. First, scorer bias is essential to address when analyzing ordinal data. The use of blinded scoring techniques, assessment, and reporting of score reliability (supported in BLISS) ameliorates this risk and quantifies reliability. Second, MIC* estimation performs best when the true MIC falls within the observed concentration range. If the estimated MIC* is higher than the highest experimental concentration tested, such results should be noted as extrapolated, as shown with triangle shapes in Figure 4. When accurate effect size estimates are essential, the concentration series should be extended until no viability is observed. Likewise, extreme floor or ceiling effects (i.e., a strain shows low or high viability at all concentrations) preclude MIC* estimation for that group, and warrant changing the range of tested concentrations. Non-monotonic treatment effects (e.g., paradoxical survival at higher concentrations) violate the assumed fixed effect of increasing concentration. In this rare case, spline-based fits are possible with PO regression (45), and theoretically with MIC* estimation, but fall outside the scope of our current framework. Lastly, a notable limitation arises when the assumption of proportional odds, which considers the effects of all explanatory variables as constant across all score categories, is strongly violated. For example, in cases with long-tail survival distributions (e.g., persister cells), the dose-response relationship can deviate from the PO model’s assumed data structure. In such cases, p-values for comparisons remain valid because the PO framework generalizes nonparametric tests, but violation of the PO assumption will lead to biased MIC* estimates, similar to any dose-response analysis of long-tailed survival data. In other words, a long tail of persisters will lead to an under-estimate of MIC*. However, a practical advantage of MIC* is that any of the aforementioned biases are easy to diagnose. When MIC* estimates fall outside of the tested concentration range, or are otherwise implausible based on observed viability across doses (such as the case of high levels of persister cells), this signals potential model violations that warrant cautious interpretation of effect size estimates. Critically, even in cases when model assumptions are violated, the PO model preserves power to detect statistically significant differences.

We envision the MIC* framework to be especially well suited to viability assays where a concentration gradient is tested using semi-quantitative scoring (e.g., plate spot assays, biofilm inhibition, or cell damage scoring). These experimental designs already generate rich, scorable phenotypes, but currently lack a standardized inferential approach. We should note that MIC* is not intended to replace standardized clinical or public health MIC breakpoints and epidemiological cutoffs, which are governed by specific standards (1, 2, 46, 47). However, for non-clinical basic and applied research, the MIC* framework enables direct estimation and hypothesis testing on absolute (ΔMIC*) or relative (Δlog_2_MIC*) scales, generating easily interpretable p-values. Extensive simulations mirroring common microbiological experimental designs, along with proof-of-principle applications, show that MIC* identifies true differences in maximum dose survived, with low bias when optimally specified, appropriate uncertainty, and desirable statistical power. Combined with BLISS for blinded scoring and ordinalMIC/MICalculator tools for analysis, this resource lowers barriers to appropriate analysis of ordinal phenotypes.

## Materials and Methods

### Yeast Case Studies

All yeast strains are listed in Table S1 and are available upon request. Our two test cases for MIC* estimation were stress survival of wild type vs mutant and a panel of different advanced intercrosses. For the wild type vs mutant comparison, cross protection data were obtained for yeast strain DBY8268 (S288C background) and an otherwise isogenic ctt1Δ mutant from (41). For the advanced intercrosses, a round-robin design (48) generated crosses of each H_2_O_2_-sensitive and resistant strain to each other. The advanced intercross was generated through 5 rounds of repeated mating and sporulation as described in (49). Yeast cross protection assays were performed as described in (38), and a detailed protocol is available at protocols.io under DOI dx.doi.org/10.17504/protocols.io.g7sbzne. Data were organized as one row per scored spot with associated covariates and sample identifiers. Spots were plated at a density of 1000 CFUs, making ≈0.1% survival the effective detection limit for MIC* estimation.

Viability at each H_2_O_2_ concentration was scored on an ordered categorical scale (lowest value indicated least viability). All images were scored in a blinded, randomized order using BLISS version 1.0.0 (Blinded Scoring System, available at LewisLabUARK.github.io/BLISS). Blinded scoring and file structure are described in detail in Supplemental Methods S1, and additional details on the BLISS software implementation can be found in Supplemental Methods S9. Acquired resistance assays were performed as described in (38), with minor modifications (Supplemental Methods), or reproduced from (41) (WT vs *ctt1Δ* mutant), with the following ordinal scoring system: high viability = 2, moderate viability = 1, and minimal or no viability = 0. The continuous predictor of interest, H_2_O_2_ concentration, was analyzed on a log-transformed scale of x=log(1+[H_2_O_2_]), to linearize the exponential effect of increasing dosage while setting zero exposure at x=0 for inclusion of unstressed control sample scores. All MIC* and ΔMIC* values are reported in the exposure scale with units of mM concentration. The intra-scorer agreement was quantified with Cohen’s weighted kappa statistic *κ* (42). Intra-scorer agreement was calculated based on repeated blinded scoring of 648 images. Discrepancies in assigned spot scores were resolved by random sampling of replicate scores.

### Proportional Odds Modelling and Definition of MIC*

Let 𝑌_𝑖_ ∈ {0, ⋯, 𝐽 − 1} be the possible ordinal viability scores for observation 𝑖, where 0 indicates “no viability”. For each 𝛼_j_ cutpoint for scores 𝑗 = 0, …, 𝐽 − 2, we fit a cumulative-logit proportional-odds (PO) model.

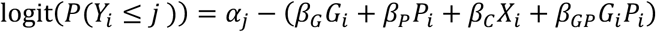

Where 𝛼_j_ are score cut-points (intercepts), 𝐺𝑖 and 𝑃𝑖 are indicator variables for genotype (e.g. WT: 𝐺_𝑖_=0, mutant: 𝐺𝑖 = 1) and pretreatment (Mock: 𝑃𝑖 =0, Pretreated: 𝑃𝑖 =1), 𝑋_i_ = log(1+[H2O2]_𝑖_), and 𝛽_𝐶_ is the slope for the effect of increasing H_2_O_2_ concentration under the proportional odds assumption. Model specification, parameterizations, and interactions follow the cumulative-logit PO framework, see Supplementary Methods S2, with extensions and alternative link functions described in Supplemental Methods S4. We use this “simplified” form (main effects plus G x P) for its inferential power and error control in observed simulations. Alternatives and when to use them are in Supplemental Methods.

For a given experimental group (𝐺 = 𝑔, 𝑃 = 𝑝), MIC* is defined as the exposure concentration at which the fitted PO model predicts 𝑃(𝑌 = 0) = 0.5, indicating “no viability” is the most probable outcome. Since the logit(0.5) = 0, MIC* is determined by the 𝑗 = 0 cumulative logit at zero, solving for 𝑋_𝐶_:

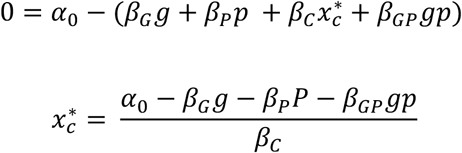

On the concentration scale (mM):

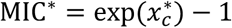

Comparison of absolute differences between experimental groups A and B are defined as ΔMIC* = MIC*_B_ - MIC*_A_, while relative comparisons can be estimated as the ratio between MIC* values, here reported on the log scale: Δlog_2_MIC* = log_2_(MIC*_B_ / MIC*_A_). Fold change in MIC* is defined as two to the power of Δlog_2_MIC*. Full algebraic derivations and closed-form gradients are in Supplementary Methods S5.

MIC* estimations provides measures of uncertainty and facilitates hypothesis testing through standard errors obtained by the multivariate delta method: the variance is 𝐺 = 𝑔^𝖳^𝛴𝑔, where 𝑔 is the gradient of the estimator (e.g., MIC* or ΔMIC*) with respect to the fitted PO model parameters (𝛼_0_, 𝛽_𝐺_, 𝛽_𝑃_, 𝛽_𝐶_, 𝛽_𝐺𝑃_) and 𝛴 is the model covariance matrix from the PO fit (derivations and expressions in Supplementary Methods S3). Wald confidence intervals are reported on estimator’s natural scale (experimental units for MIC* and ΔMIC*; log_2_ ratio for Δlog_2_MIC*), with two-sided Wald z-tests of the linear predictor for hypothesis testing of group differences. Correction for multiple testing was conducted with Benjamini-Hochberg false discovery rate correction for all a priori comparisons of interest (Supplementary Methods S17).

Diagnostics (surrogate residuals, optional Brant test) and guidance on advanced partial-PO models or mixed effect variants are provided in Supplementary Methods S8 (50). Models were fitted by maximum likelihood and retained regardless of whether the proportional odds assumption was formally met (Figure S5,S7). The connection between the proportional odds model and the nonparametric Wilcoxon rank-sum test is articulated in Supplementary Methods S7 (51). Design considerations for concentration placement around the expected MIC* are provided in SM14, with notes on numeric stability and edge-case conditions in SM15 and very small sample size options in SM16.

### Censored-Poisson Count Modelling

For Case 2, CFUs for each spot were manually counted. Spot counts above 30, or the highest number countable, were right-censored. A censored Poisson regression with the same predictor structure as the PO model was fitted using the R package VGAM version 1.1-13, and likelihood-ratio tests were used to compare coefficient estimates with those from the PO model.

### Simulation Study Design & Hypothesis Testing

To evaluate model performance, we generated synthetic datasets that reflect the yeast assay design: two genotypes, two pretreatments, twelve H_2_O_2_ concentrations, and three biological replicates (N=144). True colony counts were simulated from a Poisson distribution with

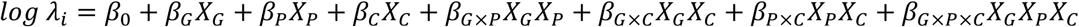

Where 𝜆_𝑖_ is the rate parameter, providing both mean and variance for cell count.

The exponentiated baseline intercept 𝑒𝑥𝑝 (𝛽_0_) corresponds to the baseline count for 𝑋_𝐺_ = 𝑋_𝑃_ = 𝑋_𝐶_ = 0. Values of 𝛽_0_ = 4.5 and Poisson model coefficients were selected to generate realistic cell counts and survival profiles across a grid of parameter values in 0.1 unit increments. Counts were binned into ordinal scores (0-4) using cut-points of 2, 10, 25, and 50 colonies. Interaction parameters were varied on a grid that produced effect sizes that ranged from 2.5 to −0.5 as change in expected count per unit change.

Comparative analysis of phenotypes based on counts used right censoring, such that counts above a threshold of 50 were censored to be >50. For each combination of parameter grid points, 1,000 datasets were simulated and analyzed with five methods: the PO ordinal model (using 𝛽 and MIC*), χ^2^ test of a contingency table, Fisher’s exact test, two-sample t-test on summed scores, and censored Poisson regression. Statistical power was calculated as the proportion of simulations with P < 0.05 when the true interaction term of interest (ΔMIC* or 𝛽_𝐺×𝑃×𝐶_) was non-zero; false positive rates (type I error) was computed with simulations where the true interaction parameter was zero. Specific model formulations tested in simulations included a full ordinal model with all two-way and three-way interactions, and a simplified model with only the genotype × pretreatment interaction and all main effects. Additional details are available in Supplementary Methods S6. Simulations were run in R v4.4.1 with a random number seed.

### Software, Data, and Code Availability

Analyses were conducted in R 4.4.1 using ordinal 2023-12-4.1 for PO model fitting and our ordinalMIC package (v1.0.0) for MIC* computation and hypothesis testing. Count-based (Poisson) models used R package VGAM 1.1-13 for censored regression. The MICalculator web app implements the identical estimator for no-code use (Supplementary Methods S10). Exact versions and session information can be found in Supplementary Materials S11. Raw plate images, scored ordinal data, and simulation scripts are be deposited in Zenodo. Source code for the MIC* calculation, multivariate delta-method variance is archived under DOI 10.5281/zenodo.17123478. Blinded scoring web application is available at github.com/LewisLabUARK/BLISS and the web tool for MIC* analysis is available for use at LewisLabUARK.github.io/MICalculator; both web tools are archived at DOI 10.5281/zenodo.17123351.

## Acknowledgments

This work was supported by the National Science Foundation grant MCB-1941824.

## Notes

https://LewisLabUARK.github.io/MICalculator

https://LewisLabUARK.github.io/BLISS

https://doi.org/10.5281/zenodo.17123478

### Competing Interest Statement

The authors have declared no competing interest.

