## Supplementary Materials for "MIC*: A Framework for Interpretable Analysis of Ordinal Viability Data"

### Supplemental Figures

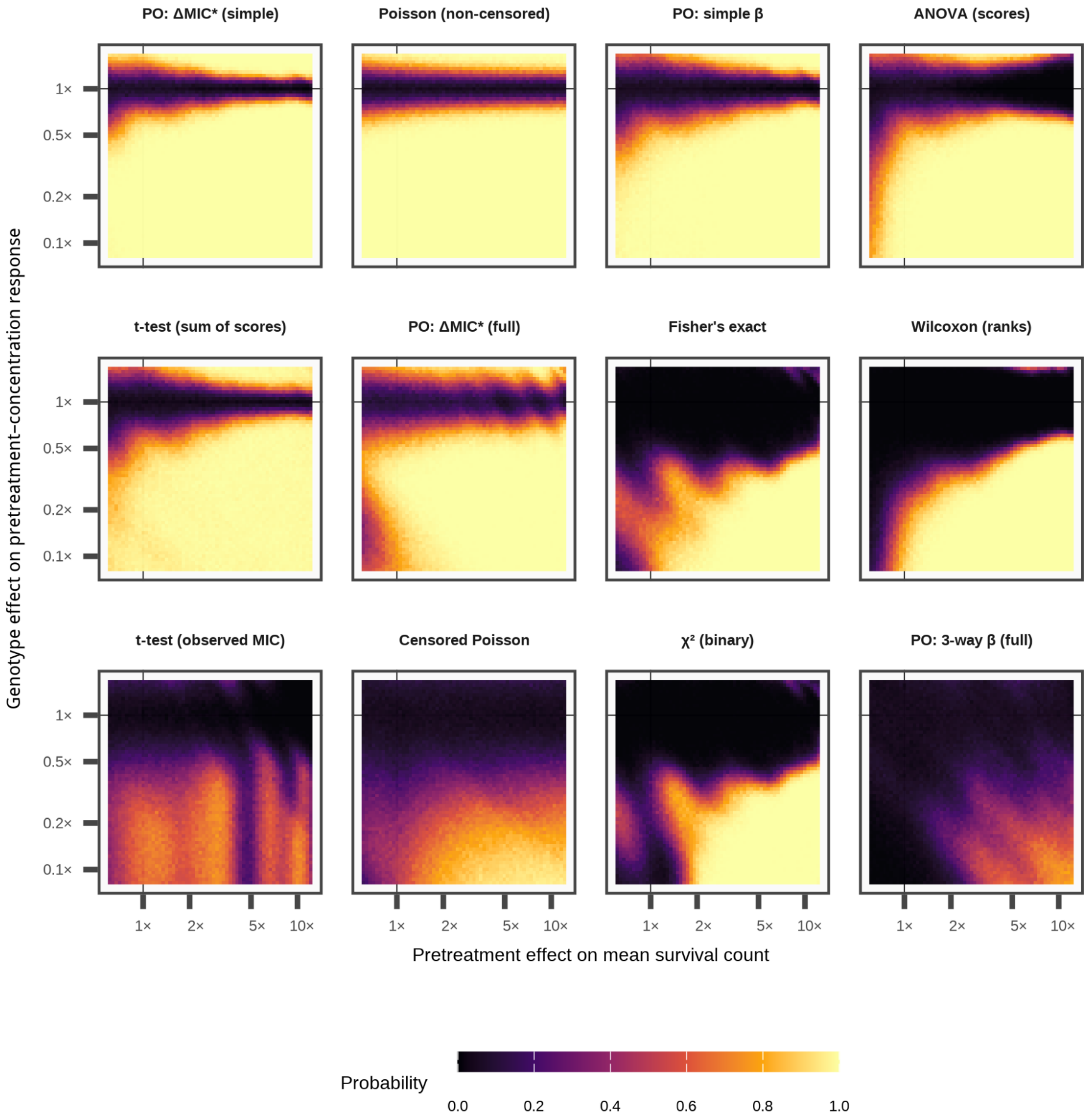


**Figure S1**. Simulation-based evaluation of statistical power (sensitivity) and Type I error (specificity, at horizontal 1x line) of each method for analyzing scored viability assays. Heatmaps show performance across a grid of treatment and genotype effect sizes. MIC* achieved the highest power while maintaining appropriate error control, with performance similar to that of the data-generating Poisson (non-censored) model. Axes show fold-change in expected survival per unit increase of log(1+concentration).

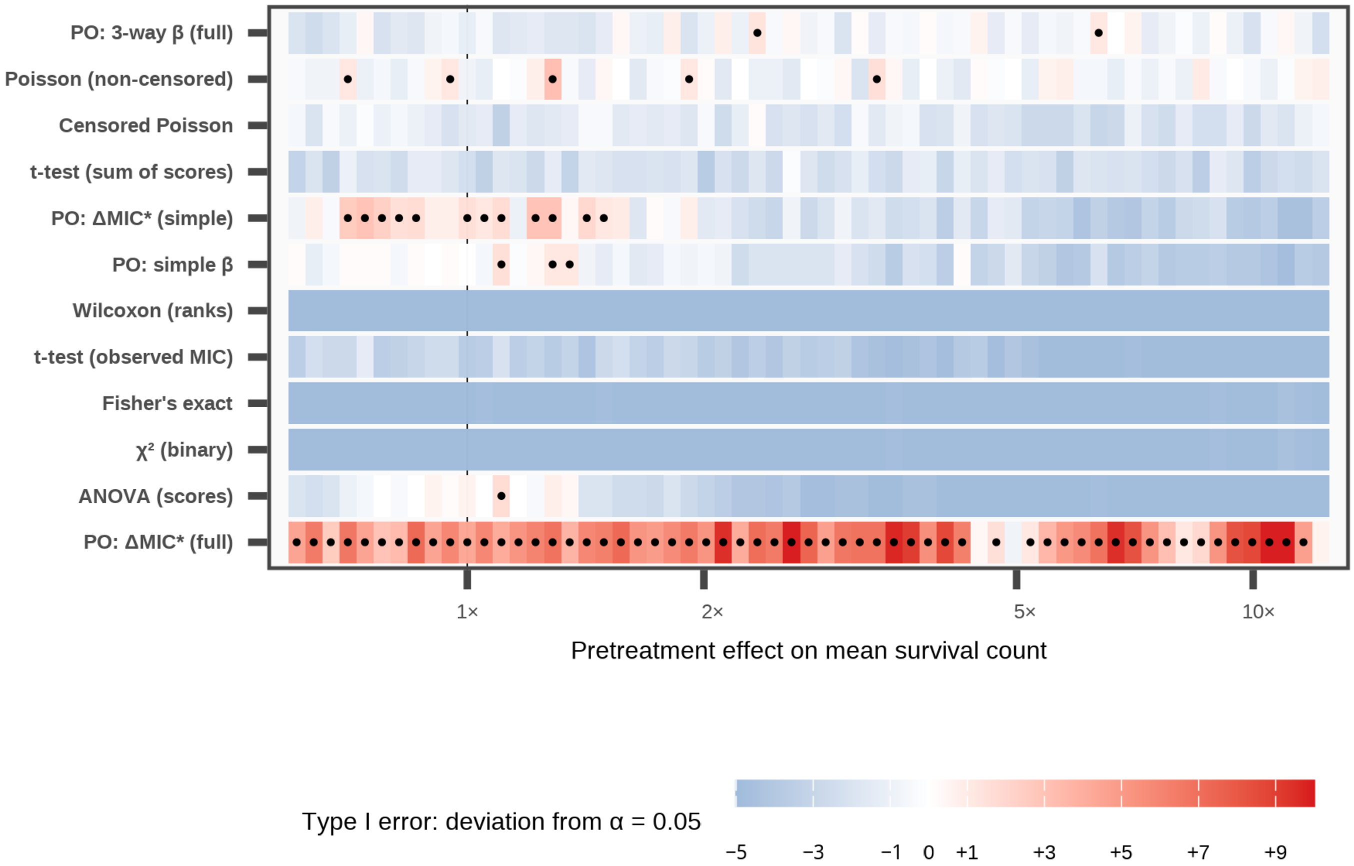


**Figure S2.** Type I error rates under the null hypothesis from simulations. Most approaches maintained appropriate or conservative error control at $\alpha = 0.05$. The saturated PO model-derived MIC* showed inflated error rates, highlighting the tradeoff between model complexity and reliability.

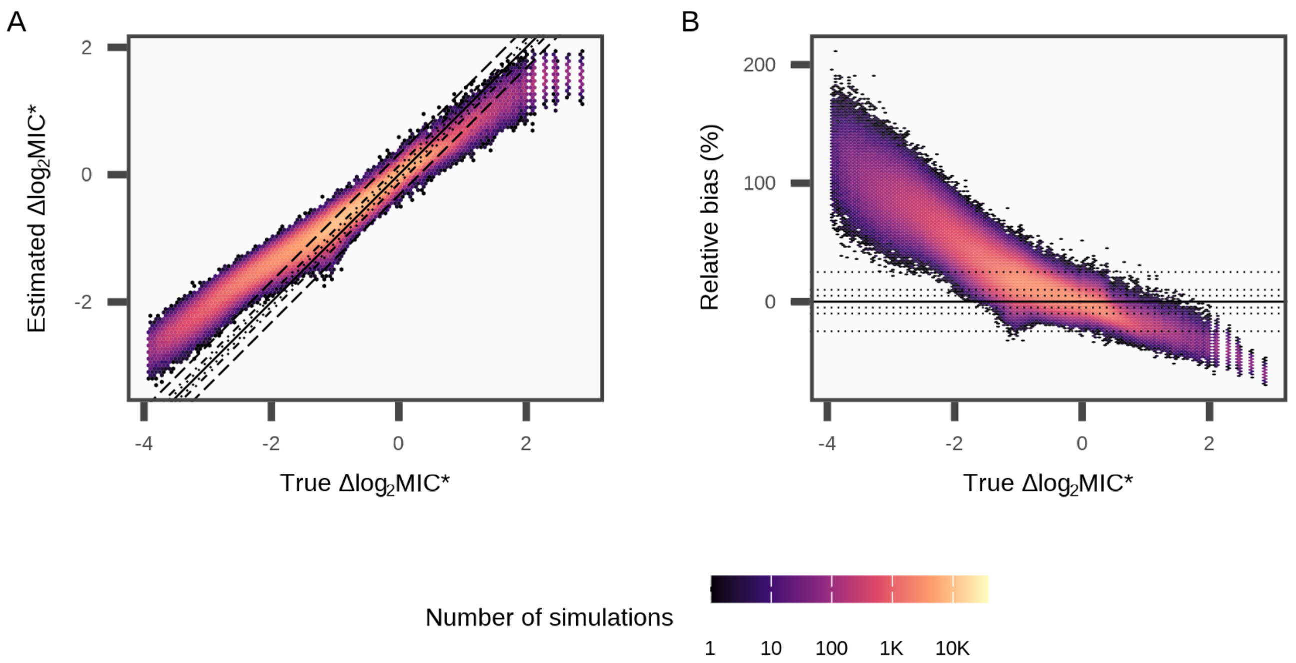


**Figure S3. (A)** Calibration and **(B)** Bias of the Simplified MIC* model as a function of true effect size. ΔMIC* estimates were slightly regularized toward smaller values, reducing overstatement of differences while preserving sensitivity. The mean relative bias for pretreated log-ratio estimates was 0.16.

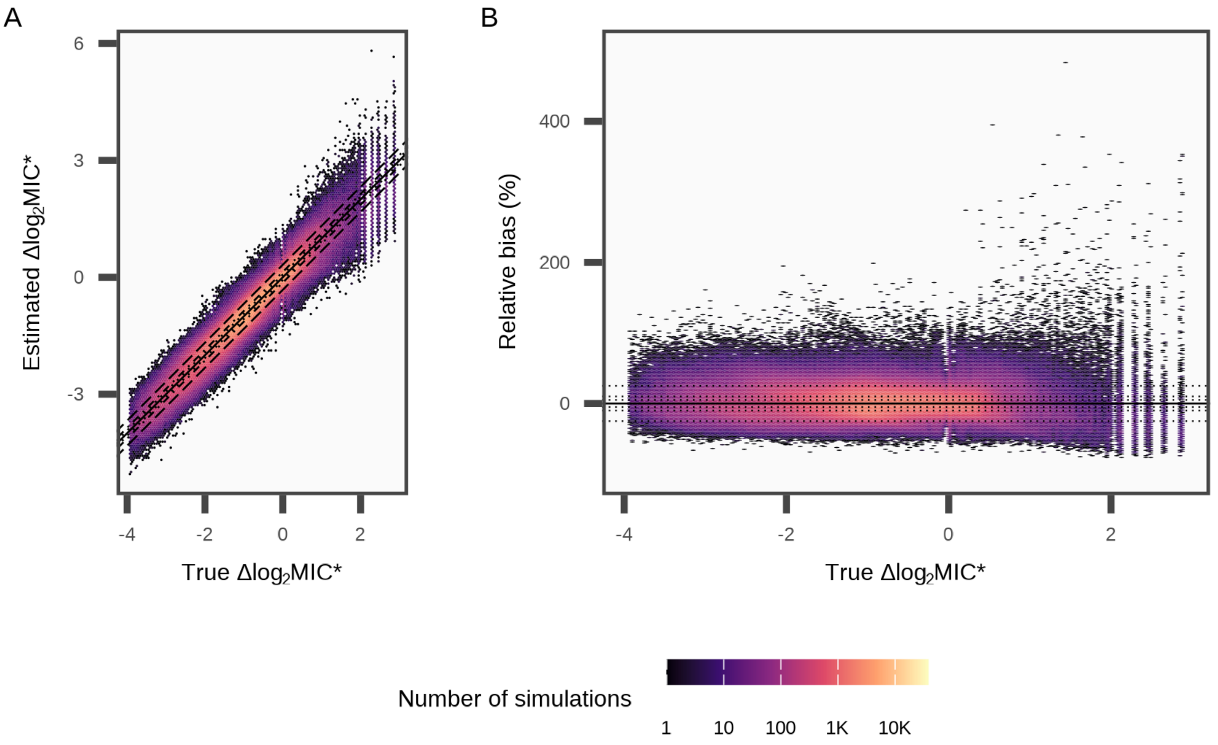


**Figure S4. (A)** Calibration and **(B)** Bias of the Saturated MIC* model as a function of true effect size. ΔMIC* estimates were unbiased on average, but showed high variability that led to unstable inference.

Six extreme outliers (>500% error) are not shown. Mean relative bias of pretreated log-ratio estimates was 0.003.

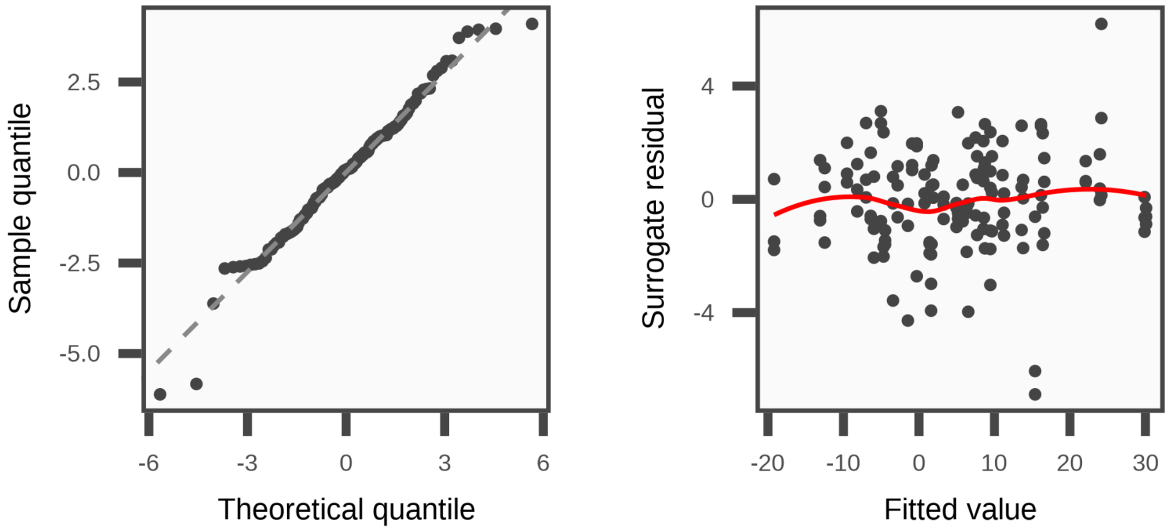


**Figure S5.** Surrogate Residual diagnostics for a representative simulated dataset. Residuals show no major deviations, supporting adequacy of the proportional-odds fit. Formal LRT-based Brant-like testing of the PO assumption identified no strong violation of the assumption (P=0.55), where P<0.05 would suggest violation of the PO assumption.

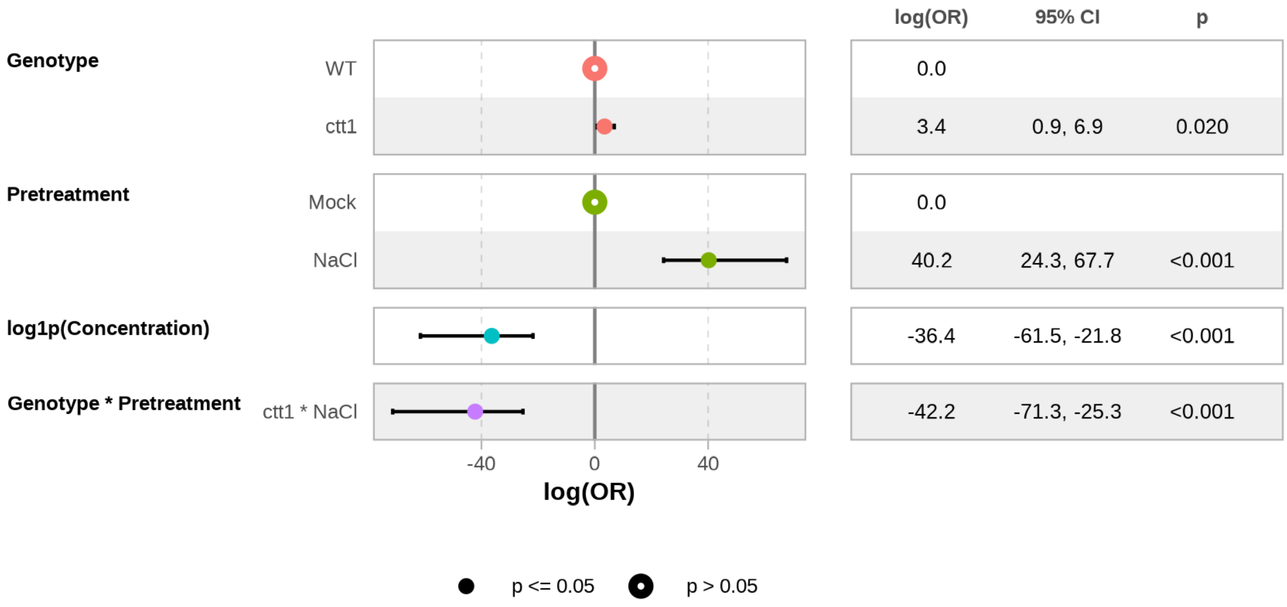


**Figure S6.** Estimated Regression coefficients for proportional-odds (PO) model for CTT1 dataset. Coefficients (log(OR)) represents the effects of strain, pretreatment, and concentration on survival scores. Baseline levels for Genotype and Pretreatment are wild-type (WT) and unstressed (Mock), respectively.

These estimates form the basis for MIC* calculation, with affect directions aligning with known conditional catalase-dependent peroxide resistance.

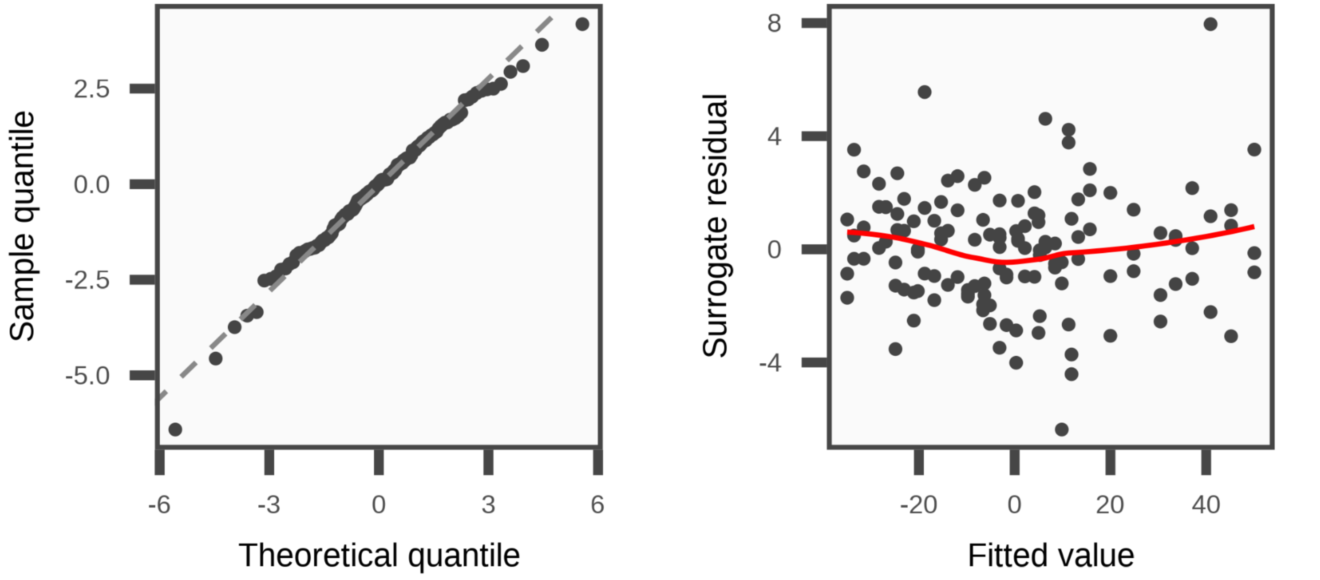


**Figure S7.** Surrogate residual diagnostics for the PO model fit to the *CTT1* dataset. Residuals are centered near zero and show no systematic trends, supporting model adequacy and appropriateness of the PO assumption in the observed data.

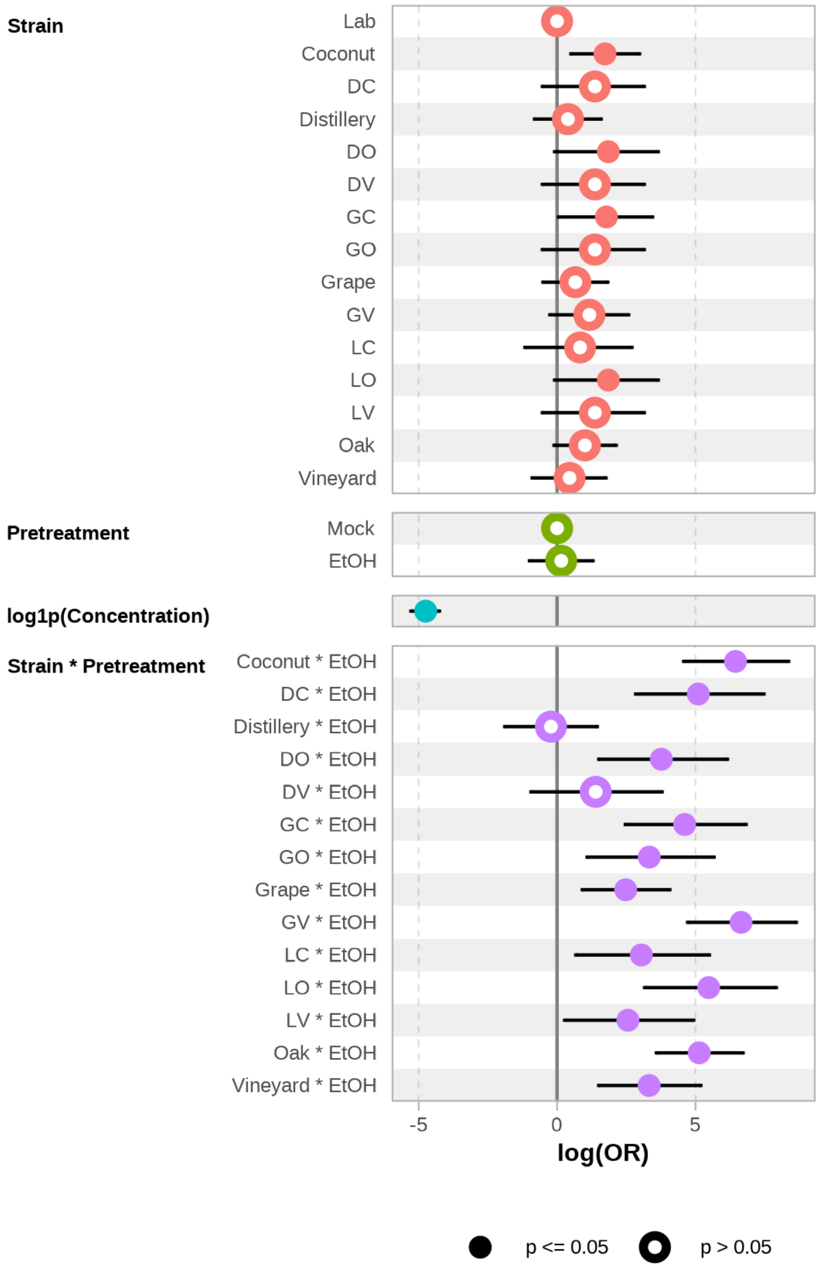


| | | log(OR) | 95% CI | p |
|---|---|---|---|---|
| Strain | Lab | 0.0 | | |
| | Coconut | 1.7 | 0.5, 3.0 | 0.006 |
| | DC | 1.4 | -0.5, 3.2 | 0.140 |
| | Distillery | 0.4 | -0.8, 1.6 | 0.521 |
| | DO | 1.9 | -0.1, 3.7 | 0.050 |
| | DV | 1.4 | -0.5, 3.2 | 0.140 |
| | GC | 1.8 | 0.1, 3.4 | 0.038 |
| | GO | 1.4 | -0.5, 3.2 | 0.140 |
| | Grape | 0.7 | -0.5, 1.8 | 0.267 |
| | GV | 1.2 | -0.3, 2.6 | 0.106 |
| | LC | 0.8 | -1.2, 2.7 | 0.392 |
| | LO | 1.9 | -0.1, 3.7 | 0.050 |
| | LV | 1.4 | -0.5, 3.2 | 0.140 |
| | Oak | 1.0 | -0.1, 2.1 | 0.078 |
| | Vineyard | 0.5 | -0.9, 1.8 | 0.505 |
| Pretreatment | Mock | 0.0 | | |
| | EtOH | 0.1 | -1.0, 1.3 | 0.797 |
| log1p(Concentration) | | -4.7 | -5.3, -4.3 | <0.001 |
| Strain * Pretreatment | Coconut * EtOH | 6.4 | 4.6, 8.4 | <0.001 |
| | DC * EtOH | 5.1 | 2.8, 7.5 | <0.001 |
| | Distillery * EtOH | -0.2 | -1.9, 1.5 | 0.795 |
| | DO * EtOH | 3.8 | 1.5, 6.2 | 0.001 |
| | DV * EtOH | 1.4 | -0.9, 3.8 | 0.245 |
| | GC * EtOH | 4.6 | 2.5, 6.8 | <0.001 |
| | GO * EtOH | 3.3 | 1.1, 5.7 | 0.004 |
| | Grape * EtOH | 2.5 | 0.9, 4.1 | 0.002 |
| | GV * EtOH | 6.6 | 4.7, 8.6 | <0.001 |
| | LC * EtOH | 3.0 | 0.7, 5.5 | 0.013 |
| | LO * EtOH | 5.5 | 3.2, 7.9 | <0.001 |
| | LV * EtOH | 2.6 | 0.3, 4.9 | 0.030 |
| | Oak * EtOH | 5.1 | 3.6, 6.7 | <0.001 |
| | Vineyard * EtOH | 3.3 | 1.5, 5.2 | <0.001 |

**Figure S8.** Estimated regression coefficients from the PO model for advanced intercross hybrid strains. Model coefficients quantify the influence of genetic background, pretreatment, and concentration. These estimates provide insights into peroxide resistance across strain combinations, but are difficult to interpret biologically. Note that strains Distillery and DV have estimated interaction effects between genotype and pretreatment not significantly different than 0. The main effect of ethanol (EtOH) describes the effect of pretreatment on the lab strain, such that the results of the PO model match the results of MIC* analysis, but requires much more expertise to accurately interpret. Furthermore, alternative formulations of the PO model can change the sign of all estimates to mean the same thing but can be very difficult to understand compared to MIC*'s clear interpretability.

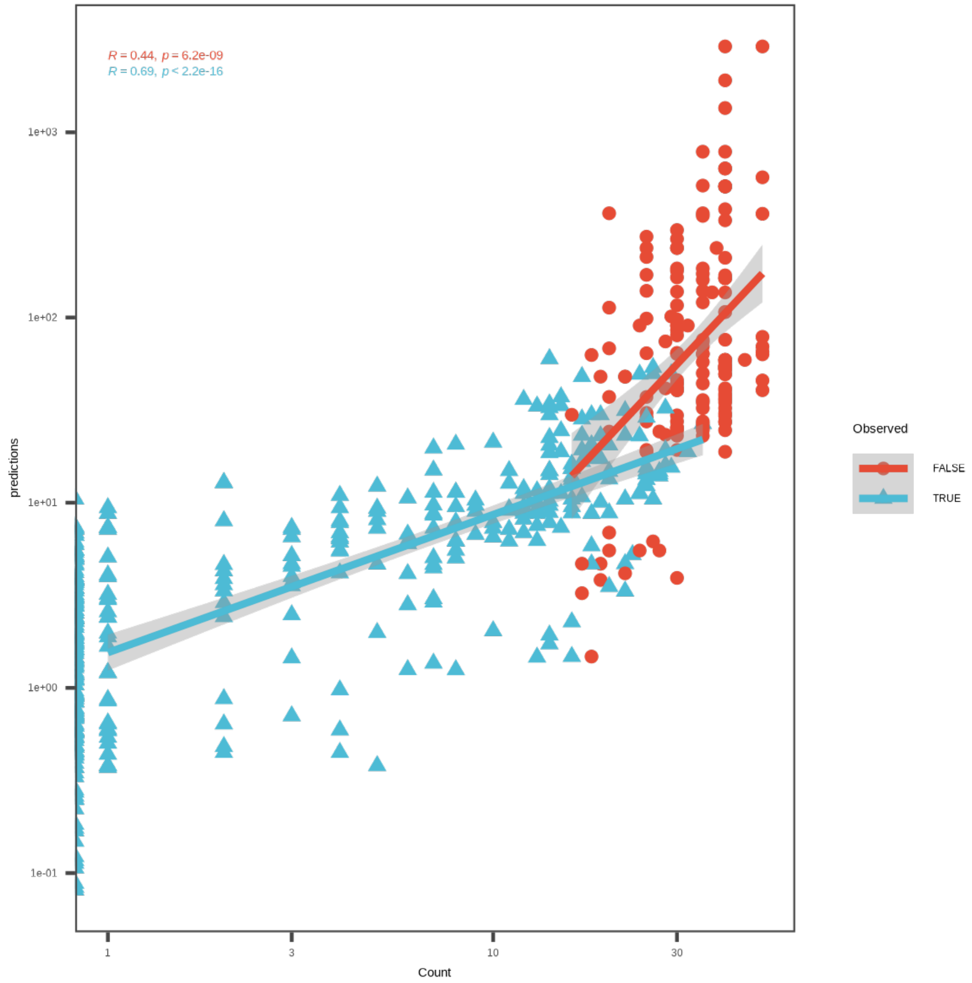


**Figure S9.** Censored colony-forming unit (CFU) estimates for each scored spot for advanced intercross hybrids. Counts were fit using a censored Poisson model with all main effects and a strain by pretreatment interaction term in order to account for colony overlaps in a spot at high colony numbers. This analysis provides a quantitative reference point for comparison with MIC* analysis of ordinal spot scores.

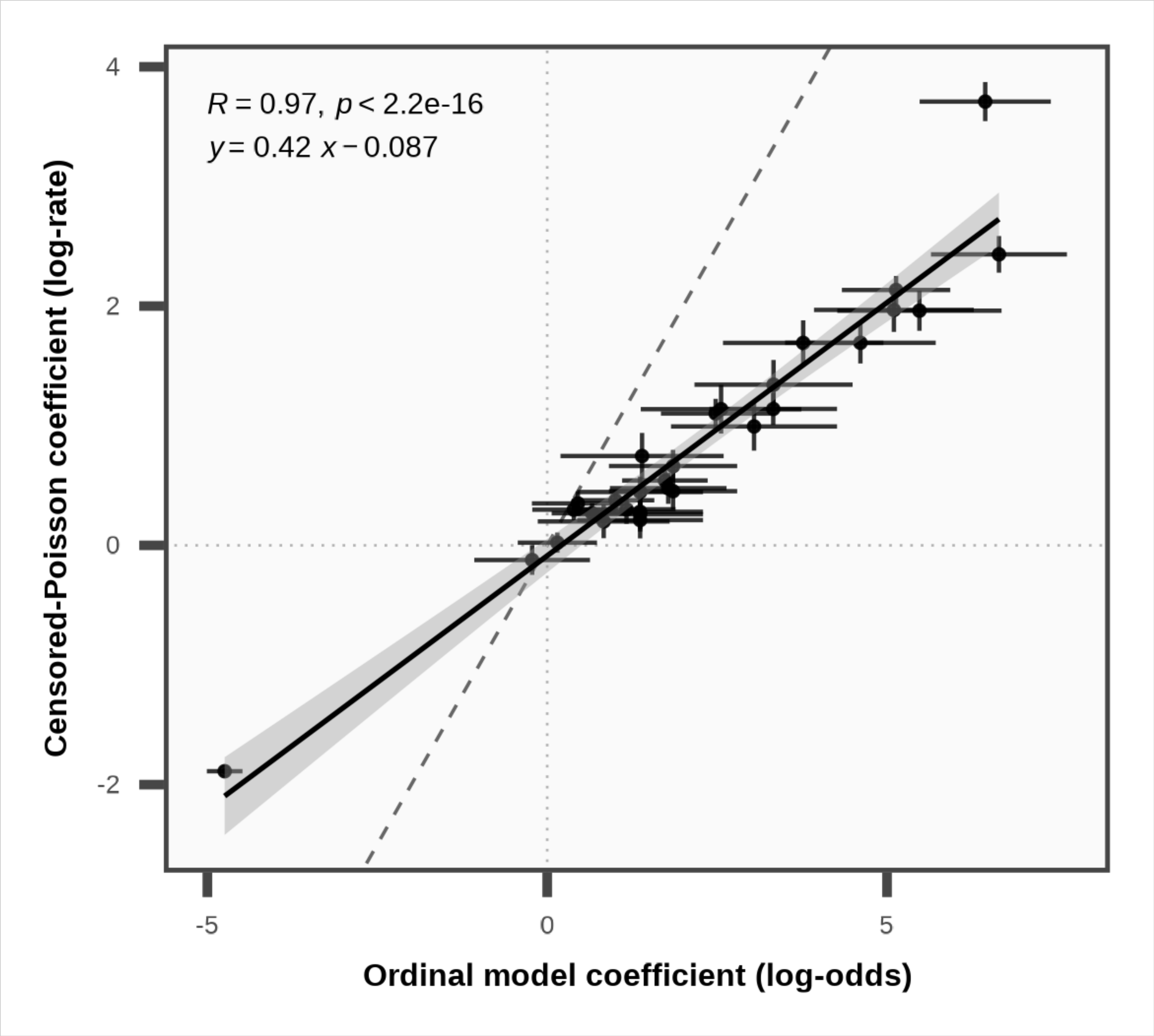


**Figure S10.** Comparison of PO and censored Poisson model coefficients. Each point represents a model coefficient estimated in both models. The strong correlation (R=0.97) indicates that PO modeling recapitulates the biological effects captured by quantitative CFU counting. While interpretations of coefficients vary between models, this result indicates similar biological meaning is being derived from ordinal spot scores and quantitative CFU counts. Error bars represent estimated coefficient standard errors.

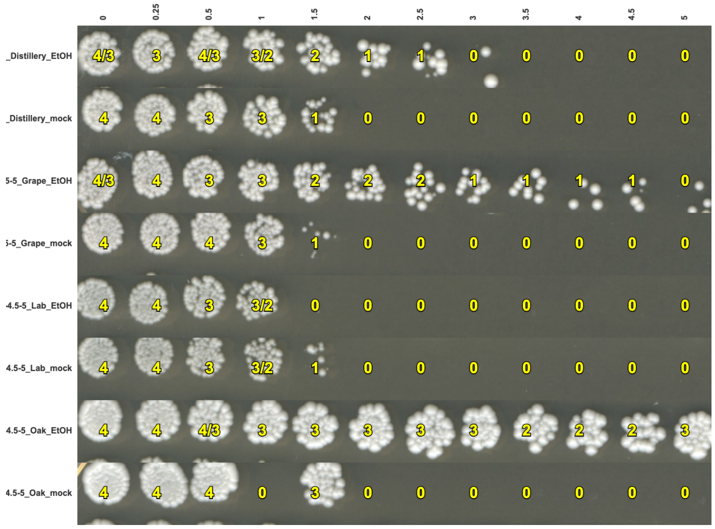


```
Scoring Summary & QC Report
-----------------------------

Images per Score Category:
- 0 (0-2 colonies): 603
- 1 (3-10 colonies): 166
- 2 (>10 colonies): 165
- 3 (TNTC): 193
- 4 (Confluent): 169
Total Images Scored: 1296

Replicate Quality Control (QC):
- Original wells with all their replicates scored: 648
- Sets of (fully scored) replicates with matching scores: 599
- Sets of (fully scored) replicates with mismatching scores: 49

Inter-Replicate Reliability (Cohen's Weighted Kappa):
  Cohen's Weighted Kappa (κ) = 0.983
  Interpretation: Almost perfect agreement
  (Calculated using Fleiss-Cohen quadratic weights, based on 648
pairs of scores from replicates of the same original images.)
```

**Figure S11.** Output from the BLISS (BLInded Scoring System) tool. Left: reconstructed image of a plate image with overlaid blindly assigned scores. Right: screenshot for the exported text report, showing scoring accuracy and reproducibility metrics. These resources enhance the utility of BLISS for rigorous and unbiased phenotype scoring in addition to providing outputs ready for MIC* analysis.

### Supplemental Tables

Table S1: Example Simulated Dataset Summarized across all tested concentrations of stressor.

| Genotype | Pretreatment | Y=0 | Y=1 | Y=2 | Y=3 | Y=4 | Total |
|---|---|---|---|---|---|---|---|
| WT | Mock | 17 | 5 | 2 | 6 | 6 | **36** |
| WT | Stress | 6 | 8 | 5 | 4 | 13 | **36** |
| KO | Mock | 18 | 5 | 1 | 6 | 6 | **36** |
| KO | Stress | 14 | 4 | 6 | 3 | 9 | **36** |
| **Total** | | **55** | **22** | **14** | **19** | **34** | **144** |

Table S2: Pairwise ΔMIC* estimates for Advanced Intercross Strain Screen.

**Table 3B. Pairwise contrasts: ΔMIC***

| Group | | Estimation | | | | | Bootstrap | |
|---|---|---|---|---|---|---|---|---|
| Group 1 | Group 2 | ΔMIC* | SE[1] | 95% CI | P | FDR (BH) | SD[1] | ΔMIC* Estimates |
| Coconut:EtOH | DC:EtOH | −7.442 | 2.508 | [-12.4, -2.53] | 0.002 | 0.003 | 0.803 | |
| Coconut:EtOH | GC:EtOH | −6.910 | 2.514 | [-11.8, -1.98] | 0.004 | 0.006 | 0.807 | |
| Coconut:EtOH | LC:EtOH | −11.643 | 2.336 | [-16.2, -7.06] | 1.87E-08 | 4.20E-08 | 0.808 | |
| Distillery:EtOH | DC:EtOH | 8.613 | 1.421 | [5.83, 11.4] | 2.32E-18 | 2.09E-17 | 0.244 | |
| Distillery:EtOH | DO:EtOH | 7.708 | 1.376 | [5.01, 10.4] | 1.68E-15 | 1.01E-14 | 0.232 | |
| Distillery:EtOH | DV:EtOH | 2.976 | 0.886 | [1.24, 4.71] | 6.49E-05 | 1.17E-04 | 0.207 | |
| Grape:EtOH | GC:EtOH | 6.257 | 1.455 | [3.41, 9.11] | 1.08E-07 | 2.17E-07 | 0.252 | |
| Grape:EtOH | GO:EtOH | 2.513 | 1.097 | [0.364, 4.66] | 0.012 | 0.015 | 0.232 | |
| Grape:EtOH | GV:EtOH | 9.359 | 1.565 | [6.29, 12.4] | 6.51E-15 | 2.93E-14 | 0.450 | |
| Lab:EtOH | LC:EtOH | 4.654 | 0.993 | [2.71, 6.6] | 1.56E-09 | 5.63E-09 | 0.227 | |
| Lab:EtOH | LO:EtOH | 11.297 | 1.860 | [7.65, 14.9] | 2.58E-23 | 4.64E-22 | 0.298 | |
| Lab:EtOH | LV:EtOH | 4.320 | 0.976 | [2.41, 6.23] | 1.61E-08 | 4.13E-08 | 0.224 | |
| Oak:EtOH | DO:EtOH | −1.092 | 1.514 | [-4.06, 1.87] | 0.482 | 0.543 | 0.401 | |
| Oak:EtOH | GO:EtOH | −3.400 | 1.254 | [-5.86, -0.943] | 0.012 | 0.015 | 0.392 | |
| Oak:EtOH | LO:EtOH | 2.254 | 1.935 | [-1.54, 6.05] | 0.217 | 0.261 | 0.441 | |
| Vineyard:EtOH | DV:EtOH | −0.705 | 1.093 | [-2.85, 1.44] | 0.523 | 0.554 | 0.148 | |
| Vineyard:EtOH | GV:EtOH | 8.567 | 1.651 | [5.33, 11.8] | 6.98E-09 | 2.09E-08 | 0.390 | |
| Vineyard:EtOH | LV:EtOH | 0.398 | 1.171 | [-1.9, 2.69] | 0.733 | 0.733 | 0.150 | |

[1] **SE (delta)**: model-based SE for ΔMIC* via delta method. **SD**: nonparametric bootstrap stratified by Strain×Pretreatment×Concentration; 1000 successful bootstraps / 0 failed refits; B = 1000.

Table S3: Pairwise $\Delta \log_2$MIC* estimates for Advanced Intercross Strain Screen.

**Table 3C. Pairwise contrasts: MIC* fold-change ($\Delta log_2 MIC*$)**

| Group 1 | Group 2 | MIC* ratio | Δlog2 MIC* | SE (log2)[†] | 95% CI | P | FDR (BH) | SD[†] | $\Delta log_2$ MIC* Estimates |
|---|---|---|---|---|---|---|---|---|---|
| Coconut:EtOH | DC:EtOH | 0.597 | −0.745 | 0.242 | [-1.22, -0.271] | 0.002 | 0.003 | 0.063 | |
| Coconut:EtOH | GC:EtOH | 0.625 | −0.677 | 0.237 | [-1.14, -0.212] | 0.004 | 0.006 | 0.063 | |
| Coconut:EtOH | LC:EtOH | 0.369 | −1.439 | 0.259 | [-1.95, -0.931] | 1.87E-08 | 4.20E-08 | 0.062 | |
| Distillery:EtOH | DC:EtOH | 4.598 | 2.201 | 0.255 | [1.7, 2.7] | 2.32E-18 | 2.09E-17 | 0.121 | |
| Distillery:EtOH | DO:EtOH | 4.220 | 2.077 | 0.262 | [1.56, 2.59] | 1.68E-15 | 1.01E-14 | 0.120 | |
| Distillery:EtOH | DV:EtOH | 2.243 | 1.165 | 0.287 | [0.602, 1.73] | 6.49E-05 | 1.17E-04 | 0.120 | |
| Grape:EtOH | GC:EtOH | 2.185 | 1.127 | 0.210 | [0.716, 1.54] | 1.08E-07 | 2.17E-07 | 0.063 | |
| Grape:EtOH | GO:EtOH | 1.476 | 0.562 | 0.222 | [0.127, 0.996] | 0.012 | 0.015 | 0.063 | |
| Grape:EtOH | GV:EtOH | 2.772 | 1.471 | 0.188 | [1.1, 1.84] | 6.51E-15 | 2.93E-14 | 0.073 | |
| Lab:EtOH | LC:EtOH | 3.163 | 1.661 | 0.274 | [1.12, 2.2] | 1.56E-09 | 5.63E-09 | 0.153 | |
| Lab:EtOH | LO:EtOH | 6.250 | 2.644 | 0.268 | [2.12, 3.17] | 2.58E-23 | 4.64E-22 | 0.154 | |
| Lab:EtOH | LV:EtOH | 3.008 | 1.589 | 0.279 | [1.04, 2.14] | 1.61E-08 | 4.13E-08 | 0.153 | |
| Oak:EtOH | DO:EtOH | 0.902 | −0.148 | 0.211 | [-0.562, 0.266] | 0.482 | 0.543 | 0.052 | |
| Oak:EtOH | GO:EtOH | 0.696 | −0.522 | 0.209 | [-0.932, -0.113] | 0.012 | 0.015 | 0.051 | |
| Oak:EtOH | LO:EtOH | 1.201 | 0.265 | 0.214 | [-0.154, 0.683] | 0.217 | 0.261 | 0.055 | |
| Vineyard:EtOH | DV:EtOH | 0.884 | −0.178 | 0.280 | [-0.726, 0.37] | 0.523 | 0.554 | 0.035 | |
| Vineyard:EtOH | GV:EtOH | 2.410 | 1.269 | 0.221 | [0.835, 1.7] | 6.98E-09 | 2.09E-08 | 0.049 | |
| Vineyard:EtOH | LV:EtOH | 1.065 | 0.091 | 0.268 | [-0.433, 0.616] | 0.733 | 0.733 | 0.036 | |

[†] **SE (log2)**: model-based SE of Δlog2 MIC* via delta method. **SD**: nonparametric bootstrap stratified by Strain×Pretreatment×Concentration; 1000 successful bootstraps / 0 failed refits; B = 1000.

### Supplemental Methods

#### S1: Data Structure and Scoring for MIC* Analysis

The MIC* framework assumes an experiment where an ordered categorical outcome (score) is observed across a series of a numeric predictor for one or more experimental groups. In our simulation and case study examples, the outcome $Y \in \{0,1,2,3,4\}$ is an ordinal viability score recorded for each spotted culture at pre-specified concentrations of stressor. The numeric predictor is the concentration of hydrogen peroxide stressor reported in its natural units (mM), optionally transformed for modeling. Each observation corresponds to a single scored "spot" (or well), accompanied by its grouping covariates (e.g., genotype, treatment, or strain) and, when applicable, replicate identifiers. We used custom tools BLISS and ordinalMIC, described in S9-10 below, for all MIC* analysis described.

#### S2: Proportional-Odds (PO) Model Specifications

Let $Y \in \{0,1,\ldots,J\}$ denote the ordinal outcome with the lowest score "0" representing inhibition. Let $x_c$ be the numeric variable of interest used in the model for the experiment (e.g., $x_c = \log(1 + \text{mM H}_2\text{O}_2]))$, and let $x_{nc}$ be a vector of other covariates (e.g., genotype, pretreatment, etc.). The cumulative-logit PO model is

$$log\left(\frac{P(Y \le j)}{P(Y > j)}\right) = \alpha_j - \eta(X), j = \{0,\ldots,J-1\}$$

with cut points $\alpha_0 \le \alpha_1 \le \ldots \le \alpha_{J-1}$, and linear predictor $\eta(X) = \eta(x_c) + \eta(x_{nc})$. Interaction terms can be included in this type of model, including both within $x_{nc}$ and between $x_c$ and $x_{nc}$ variables included in the model. This parameterization follows the ordinal package in R, in which positive $\beta$ coefficients correspond to higher probability of higher scores, and vice versa. Models were fit by maximum likelihood with logit (i.e., $\log\left(\frac{Z}{1-Z}\right)$) link function and Hessian-based covariance estimates.

#### S3: MIC* Defined on Experimental Scale

We define MIC* for covariates setting $x_{nc}$ as the concentration $c^*$ at which the fitted model predicts $P(Y = 0|x_c^*, x_{nc}) = 0.5$. Because $P(Y = 0) = P(Y \le 0)$ as 0 is the lowest score, and $logit(0.5) = log\left(\frac{0.5}{1-0.5}\right) = log(1) = 0$, solving for MIC* is equivalent to solving

$$\alpha_0 - \eta(x_c^*, x_{nc}) = 0.$$

For the predictor $x_c^*$ on the modeling scale and then mapping back to the experimental scale by $c^* = f^{-1}(x_c^*)$ where $f(c) = log(1 + c)$ in examples shown. Under the simplified linear form $\eta(x_c, x_{nc}) = \beta_c x_c + x_{nc}^\top \beta_{nc}$, the solution is explicit:

$$x_c^*(x_{nc}) = \frac{\alpha_o - x_{nc}^\top \beta_{nc}}{\beta_c}$$

$$MIC^*(x_{nc}) = exp\,\{x_c^*(x_{nc})\} - 1$$

The experimental and modeling scale are distinguished for the calculation process, such that on the experimental scale the concentration is denoted as $c$ (e.g., mM $H_2O_2$) while on the modeling scale the predictor $x_c = f(c)$ is used for the transformed dose value for linearizing dose-response effects (e.g., $x_c = log(1+c)$). When no transformation is applied to the $x_c$ predictor, $c = x_c$. For clarity, this notation is omitted from the main text, in which concentration is only referred to on the experimental scale.

Note the linear predictor is denoted as $\eta(x_c, x_{nc})$, where $x_{nc}$ collects all grouping (non-concentration) covariates (e.g., genotype, pretreatment). In the main text, these effects are defined more simply as factors $G$ (genotype), $P$ (pretreatment), $C$ (concentration effect), and $GP$ (interaction). Both are equivalent descriptions of the PO model.

Uncertainty is obtained analytically via the multivariate delta method. Let $\theta$ be a vector of model parameters $\alpha_0, \beta_c, \beta_{nc}$ and let $\Sigma = Cov(\hat{\theta})$. Define $g(\theta) = MIC^*(x_{nc})$ as above. Then,

$$Var\{\widehat{\text{MIC}}^*(x_{nc})\} \approx \nabla g(\hat{\theta})^\top \sum \nabla g(\hat{\theta})$$

with gradient components derived in Supplemental Methods 5. Pairwise differences are handled accordingly, with $\Delta MIC^* = MIC_B^* - MIC_A^*$, and $\Delta \log_2 MIC^* = \log_2 MIC_B^* - \log_2 MIC_A^*$ using the same $\Sigma$ matrix as both endpoint estimates are functions of a single fitted PO model.

### S4 Alternative Link Functions and Scale Shift Parameters

We use the logit link by default, but MIC* analysis can be extended directly to other cumulative link functions. For any link function $g(\cdot)$, with inverse $g^{-1}(\cdot)$, the PO model can be written as $g\{\, P(Y \le j)\,\} = (\alpha_j - \eta)/s$ where $s > 0$ is an optional scale parameter, as formulated in the ordinal R package's location-scale extension. Setting $P(Y \le 0) = 0.5$ yields

$$\alpha_0 - \eta(x_c^*, x_{nc}) = sC_{link}, \quad C_{link} \equiv g(0.5)$$

The linear predictor $x_c^*$ for the experimental group $x_{nc}$ can be calculated as

$$x_c^*(x_{nc}) = \frac{\alpha_0 - x_{nc}^\top \beta_{nc} - sC_{link}}{\beta_c}$$

For symmetric links (logit and probit) with $C_{link} = 0$, the scale $s$ cancels, meaning the MIC* estimates are invariant to scale parameters. For asymmetric link functions log-log and complementary log-log, a scale-parameter sensitive fixed offset is required for MIC* estimation, with $C_{cloglog} = \log(-\log(1-0.5)) = \log(\log(2)) \approx -0.367$ for the complementary log-log link, while for the log-log link, $C_{loglog} = -\log(-\log(1-0.5)) = -\log(\log(2)) \approx 0.367$. Note these constants are multiplied by the fitted scale parameter $s$ if a scale model is used, by default $s = 1$. After computing $x_c^*$, the experimental-scale MIC* is again $\text{MIC}^* = exp(x_c^*) - 1$ for the natural log transformation with pseudocount of one we use here, or in general the $\text{MIC}^* = f^{-1}(x_c^*)$ for transformation $f(\cdot)$ applied to the natural numeric scale.

### S5 Complete Derivations of MIC*, ΔMIC*, and Δ$\log_2$MIC*

Let $f(c) = log\,(1+c)$ and $f^{-1}(x) = exp(x) - 1$. Define

$$T(\boldsymbol{\theta}) \equiv \frac{\alpha_0 - x_{nc}^\top \beta_{nc} - sC_{link}}{\beta_c}, \text{ and } c^*(\boldsymbol{\theta}) \equiv f^{-1}\{T(\boldsymbol{\theta})\}.$$

Since $f'(c) = \frac{1}{1+c}$, the chain rule gives

$$\frac{\partial c^*}{\partial T} = \frac{1}{f'(c^*)} = 1 + c^*$$

Hence the gradient components of $c^*$ are

$$\frac{\partial c^*}{\partial \alpha_0} = \frac{1 + c^*}{\beta_c}, \quad \frac{\partial c^*}{\partial \beta_c} = -(1 + c^*)\frac{T}{\beta_c}, \quad \frac{\partial c^*}{\partial \beta_k} = -(1 + c^*)\frac{x_k}{\beta_c},$$

for any coefficient $\beta_k$ in $\mathbf{x}_{nc}^{\top}\boldsymbol{\beta}_{nc} = \sum_k x_k \beta_k$. For example, $x_k = 1$ for the presence of indicator variable $k$ or can equal the covariate's value for a numeric model term. In the case of a scale model being used, $\partial c^*/\partial s = -(1 + c^*)C_{link}/\beta_c$. These expressions, evaluated at $\widehat{\boldsymbol{\theta}}$, form the gradient $\nabla c^*$ used in calculation of $Var(\widehat{c^*}) \approx \nabla^{\top}\Sigma\nabla$.

For $\Delta MIC^* = c_B^* - c_A^*$, defined $g_{\Delta}(\boldsymbol{\theta}) = c_B^*(\boldsymbol{\theta}) - c_A^*(\boldsymbol{\theta})$. The gradient is the difference of individual gradients, evaluated at the corresponding covariate encodings. For $\Delta \log_2 MIC^* = \log_2 c_B^* - \log_2 c_A^*$, application of the chain rule provides

$$\frac{\partial}{\partial \theta} log_2 c^* = \frac{1}{c^* \, log\,(2)} \cdot \frac{\partial c^*}{\partial \theta}$$

Such that the gradient is $\nabla \log_2 c_B^* - \nabla \log_2 c_A^*$. Wald $Z$ statistics are calculated from these variances directly.

When the linear predictor contains interactions with $x_c$, (e.g., $x_c \times G$, $x_c \times P$, or $x_c \times G \times P$), the solution for $x_c^*$ is still closed form as $\eta$ is still linear in $x_c$. For example, in the saturated model where $\eta = \beta_c x_c + \beta_g G + \beta_p P + \beta_{gc}(Gx_c) + \beta_{pc}(Px_c) + \beta_{gp}(GP) + \beta_{gpc}(GPx_c)$, let all terms not containing $x_c$ into $A = \beta_g G + \beta_p P + \beta_{gp}(GP)$ and all terms containing $x_c$ into $B = \beta_c x_c + \beta_{gc}(Gx_c) + \beta_{pc}(Px_c) + \beta_{gpc}(GPx_c)$. It follows that

$$x_c^* = \frac{\alpha_0 - A - sC_{link}}{B}, \qquad c^* = exp(x_c^*) - 1,$$

with gradients obtained by differentiating w.r.t. $A$ and $B$.

### S6 Simulation-Based Estimation of Power and Effect Sizes

The simulation study was implemented in R to mirror routine microbiological designs, in which an ordered viability score is recorded across a fixed series of concentrations for two genotypes, each tested with and without some pretreatment. For each simulation ran, a dataset comprised of 2 by 2 experimental groups (Genotype $G \in \{a, b\}$ and Pretreatment $P \in \{no, yes\}$, a grid of 12 pre-defined stressor concentrations, and three biological replicates per group, yielding $N = 2 \times 2 \times 12 \times 3 = 144$ observations to match biological sample size and layout. The concentrations series used in simulations were $\{0,1,2,2.5,3,3.5,4,5,7,10,25,50\}$, transformed for modeling on the $x_c = \log(1 + \text{concentration})$ scale such that zero exposure maps to $x_c = 0$ and accounting for the quasi-multiplicative effect pattern of original units. Genotype and pretreatment covariates were coded as binary indicators $G \in \{0,1\}$ and $P \in \{0,1\}$ with genotype "A" and "no" pretreatment as the corresponding baseline values.

Data were generated at the level of colony counts using a Poisson GLM model of the form:

$$\log(\lambda(G, P, x_c)) = \beta_0 + \beta_G G + \beta_P P + \beta_C x_c + \beta_{GP} GP + \beta_{GC} G x_c + \beta_{PC} P x_c + \beta_{GPC} GP x_c$$

Such that the expected count at some combination $(G, P, x_c)$ equals $\lambda = exp(\cdot)$. The coefficient $\beta_C < 0$ produces decreasing viability with increasing stressor concentration, while $\beta_{PC}$ determines the effect of pretreatment on survival under non-zero stressor concentrations and $\beta_{GPC}$ controls the extent to which the alternate genotype affects survival at increasing concentrations when pretreated. Baseline cell counts without stress exposure were set to be 1000 by $\beta_0 = 6.91$. The effect of genotype $\beta_G = 0$ to emulate equal survival when unstressed, the main effect of pretreatment ($\beta_P = 0$) as pretreatment had no effect on cell survival when exposed to no stressor. Both $\beta_{GP} = \beta_{GC} = 0$ to match the assumption of pretreatment having no different effect on unstressed survival between genotypes and genotype having no effect on basal resistance to stressor, respectively. The final two parameters were simulated at different values to reflect realistic possible experimental outcomes, with effect of pretreatment on stress survival $\beta_{PC} \in \{-0.5,2.5\}$ and effect of genotype on reduced stress survival following pretreatment $\beta_{GPC} \in \{-2.5,0.5\}$, each tested over a grid of 0.1 increments for a total of $31^2 = 961$ parameter combinations, each simulated 1000 times using Monte Carlo simulations for a total of $9.61\times10^6$ simulated experiments. Realized counts $Y_{count} \sim Pois(\lambda)$ were then mapped to the scale of ordinal scores $Y \in \{0,1,2,3,4\}$ using fixed cut-points (2,10,25,50) such that $Y = 0$ for $Y_{\text{count}} \leq 2$, $Y = 1$ for 3 to 10, $Y = 2$ for 11 to 25, $Y = 3$ for 26 to 50, and $Y = 4$ for $Y_{\text{count}} > 50$. This mapping approximates common spot assay scoring and creates a realistic mixture of score distributions across the concentration gradient.

Each simulated dataset was analyzed by two proportional-odds (PO) cumulative-logit model specifications, using our **ordinalMIC** package: a "Saturated" model including all two-way and three-way interactions for $(G, P, x_c)$, and a "Simplified", more parsimonious model with all main effects for $(G, P, x_c)$, and a single two-way interaction for $G \times P$ to estimate the effect of genotype on pretreatment, with a common affect fit for $x_c$. From each fitted PO model in each simulation, MIC* estimates were obtained for the four experimental groups: $(G,P) \in \{(0,0),(0,1),(1,0),(1,1)\}$ by solving for $P(Y \leq 0 \mid G, P, x_c^*) = 0.5$ for $x_c^*$ and back-transforming the concentration with $\text{MIC}^* = \exp(x_c^*) - 1$. Pairwise differences between groups were computed on both the absolute scale, $\Delta\text{MIC}^* = \text{MIC}_B^* - \text{MIC}_A^*$, and on the relative scale, $\Delta \log_2 \text{MIC}^* = \log_2(\text{MIC}_B^* / \text{MIC}_A^*)$, using the multivariate delta method and the model's Hessian matrix. The implementation utilized the mic_solve() function in the **ordinalMIC** package to return group-wise MIC* estimates with analytic standard errors, contrast tables for $\Delta MIC^*$and $\Delta \log_2 MIC^*$ alongside Wald Z tests and confidence intervals. A difference of differences estimate is also provided on relative and absolute scales to summarize $G \times P$ effects.

For comparing performance of the MIC* estimator against common alternatives, we fit several additional models and applied statistical tests to the same simulated datasets. The same Poisson GLM model used to generate the data was also fit to the resulting counts directly (rather than the corresponding scores) to serve as a benchmark analysis using the optimal quantitative analysis on the count scale. Because in practice true CFUs of spots cannot be observed due to crowding, we fit a right-censored Poisson regression (cens.poisson family in the VGAM R package v1.1-13) in which counts above 50 were right-censored to approximate practical counting limits while maintaining the same model function. Relevant model coefficients were tested by likelihood-ratio tests.

Alternative approaches to hypothesis testing for group differences are frequently applied to ordinal data. Nominal tests are often used to analyze ordinal data across concentration series, which we compared by applying $\chi^2$ and Fisher's Exact tests to the 2 by 5 genotype-by-score table among observations where $P = 1$. Additionally, we applied the Wilcoxon rank-sum test to compare scores between genotypes at $P = 1$, where concentration is not modeled. A two-sample t-test with unequal variance was applied to (i) the observed MIC (smalled concentration where $Y = 0$ observed) for triplicate replicates, and (ii) the per-replicate sums of scores of pretreated samples between genotypes. A two-way ANOVA treating score as a numeric value using genotype, pretreatment and concentration as categorical factors, and extracting the $G \times P$ term to parallel the simplified PO contrast. An auxiliary ordinal model that treated the raw counts as ordered categories, rather than binned into scores, was also fit to explore the effect of varying the number of score categories.

We summarized the magnitude of the $\beta_{GPc}$ interaction as the per-unit change in expected counts under pretreatment in the mutant relative to wild type across the unit of concentration, such that reduction in survival is calculated as $1 - \exp(\beta_{GPc})$. For example, $\beta_{GPc} = -0.36$ corresponds to $\approx 30.3\%$ reduction per unit of $x = log(1 + c)$. This definition aligns the Poisson generating model with the sign and magnitude conventions used when plotting power vs percentage reduction.

The "true" MIC* in simulations was computed directly for each group from the data generating model and reported on the experimental scale. Using the chosen category cutoffs, we solved for $P(Y \leq 0) = 0.5$ under the Poisson distribution. Let $\lambda_{50}$ denote the Poisson rate parameter at which $P(Y_{\text{count}} \leq 2) = 0.5$. Solving for $F(2; \lambda_{50}) = 0.5$ provides $\lambda_{50} \approx 2.674$. Let $log\ \lambda = A + Bx_c$ for group-specific $A$ and $B$, the theoretical MIC is then:

$$x_c^{\dagger} = \frac{log\ \lambda_{50} - A}{B}, \qquad \text{True MIC} = \exp\left(x_c^{\dagger}\right) - 1$$

Where $\text{True MIC} = \infty$ when $B \geq 0$, corresponding to no killing effect of concentration. Group-wise quantities for $A$ and $B$ for each $(G, P)$ are calculated using model parameters for the data generating function of each simulation.

The simulation function returned, for each replicate and parameter combination, MIC* point estimates for all four groups, Wald test p-values and confidence intervals for ∆MIC* and $\Delta\log_2$MIC* for both simplified and saturated PO models, *p*-values for the interaction term in PO models directly, and *p*-values for all procedures used for comparison. After aggregating results across simulation replicates, performance metrics of each approach were computed in the following manner. Simulations in which $\beta_{GPC} = 0$ were designated as null for type I error rate estimation at $\alpha = 0.05$, while all other scenarios were analyzed as possessing some true nonzero ∆MIC* and were tested for power. Bias and root-mean-squared error (RMSE) for MIC* were obtained by comparing estimated MIC* to the corresponding True MIC derived from the data generating process. Because all MIC* contrasts are smooth functions of a single PO fit, uncertainty summaries and joint inference procedures (e.g., familywise error control across multiple contrast) were derived from the same delta method-based covariance. Details of multiplicity adjustments and small-sample considerations are discussed in supplemental methods sections S16-17.

### S7 Wilcoxon Rank-Sum Test Connection to the PO Model

The PO model provides a regression generalization of the Wilcoxon rank-sum (Mann-Whitney U) test. Consider the two-sample case with binary group indicator $G \in \{0,1\}$ and ordinal outcome $Y \in \{0,1,\ldots,J-1\}$. The proportional odds (PO) cumulative link model with an arbitrary set of category cutpoints $\alpha_j$ specifies

$$\text{logit}\{P(Y \leq j|G)\} = \alpha_j - \beta G, \qquad j = 0,\ldots,J-2,$$

With $\beta$ the common log-odds ratio parameter. The null hypothesis $H_0 : \beta = 0$ states that the two cumulative distributions coincide at every score threshold, meaning no global shift is present in the distribution of $Y$ between groups.
In this setting, the efficient score test of $H_0 : \beta = 0$ from the PO likelihood is asymptotically equivalent to the Wilcoxon/Man-Whitney (WMW) two-sample rank-sum test (with midranks), up to a constant scale factor. This can be seen by writing the PO log-likelihood $\ell(\alpha_j, \beta)$, differentiate with respect to $\beta$, and evaluate the score at $\beta = 0$ after profiling out $\alpha_j$. The resulting score statistic can be expressed as a sum over categories of group-centered indicators $\mathbf{1}\{Y \le j\}$, weighted by the estimated marginal probabilities $\hat{\pi}_j = P(Y \le j)$ under $H_0$:

$$U_\beta \big|_{\beta=0} = \sum_{i=1}^{n} (G_i - \bar{G}) \sum_{j=0}^{J-2} \frac{\mathbf{1}(Y_i \le j) - \hat{\pi}_j}{\hat{\pi}_j(1 - \hat{\pi}_j)}$$

Note that, after algebraic simplification, the inner summation is a strictly increasing function of the pooled midrank of $Y_i$. Hence, the score test reduces to a linear rank test with Wilcoxon scores (midranks), which is the WMW test. Because the nuisance cutpoints $\alpha_j$ are estimated under $H_0$, and only enter through $\hat{\pi}_j$, the equivalent holds without requiring distributional assumptions beyond ordinality. Thus, for two groups and no covariates, testing $\beta = 0$ in the PO model is the Wilcoxon test, up to scaling, and the PO model may be regarded as a covariate-generalization of the nonparametric WMW. Formal statements and proofs are given in McCullagh 1980, Agresti 2010, and Harrell 2015, in which the rank-test equivalence via score functions (or the ridit/concordance formulations) are derived.

Two immediate corollaries are important in practice. First, because WMW is a valid nonparametric test under broad conditions, mild violations of the PO assumption primarily impact efficiency (i.e., power and standard errors), not the validity of the group comparison. This connection to the PO model drives our perspective PO model testing and diagnostics (e.g., surrogate residuals). Second, the PO coefficient $\beta$ is a monotone function of the probability of stochastic superiority, $p = P(Y_1 > Y_0) + \frac{1}{2}P(Y_1 = Y_0)$, which is the estimand targeted by the Wilcoxon rank-sum test. Indeed, $p$ relates directly to concordance or Somer's $D$ with $D = 2p - 1$. This means that direction of affect interpretations align between WMW and the PO model as $sign(\beta) = sign(p - 0.5)$.

This connection extends naturally to when additional covariates enter the PO model structure. Whether additional factors or continuous predictors (e.g., transformed exposure $x = log(1 + c)$ used to define MIC*), the PO model retains the rank-based approach of WMW while enabling adjusted and interaction containing inference. MIC* leverages this structure by converting the fitted PO surface into an exposure-scale quantity. This preserves robustness and interpretability of WMW-type inference while placing estimation and hypothesis testing on the experimental scale via delta method approximation.

### S8 Diagnostics and Model Construction

The adequacy of model fits was examined using surrogate residuals for ordinal regression, which under the PO assumption exhibit mean near zero and no structure against fitted values or predictors. A brant-type check of proportional odds can be computed but note that interpretations in the MIC* context derives from concentration-scale contrasts. Violations of the PO assumption influences estimator bias and precision, but the validity of comparisons is conserved. Given strong biological reasoning to suspect non-parallel effects of covariates at specific score categories, partial PO models facilitate coefficients to vary by cutpoint.

Notably, the definition of MIC* allows for estimation with partial- or non-proportional odds models, but power can suffer in this context as the amount of data utilized is reduced. Similarly, hierarchical designs and repeated observations (e.g., time series of images for a plate) can be accommodated with random-effects cumulative link models (implemented in ordinal R package with clmm function). The delta-method formulas for estimation can use the $\Sigma$ matrix from a mixed-model fit. For non-monotone phenotypes, such as paradoxical survival at higher concentrations, spline terms in $x_c$ can theoretically be introduced, although this is beyond the scope of this manuscript. MIC* would then be interpreted as a local crossing point, where we recommend mapping uncertainty by simulation rather than relying on first-order approximations.

### S9 BLISS Implementation Details

BLISS is a browser-based tool for image preparation and blinded scoring that runs entirely on the user's local machine with no installation required. We host this tool (MIT license) online for easy access with an internet connection at LewisLabUARK.github.io/BLISS. Alternatively, the html file can be downloaded from https://zenodo.org/records/17123352 for fully offline use. Consisting of two main components, BLISS provides a tool for image segmentation (Image Dicer) and Scoring App for unbiased score assignment.

Image Dicer uses HTML5 Canvas to segment a grid (plate) of observations (wells) contained in an image into an array of images of individual observations (wells). The image-segmenting grid is applied based on user-defined row and column count while relevant labels for rows and columns can be added for downstream analysis. The Image Dicer crops each image into a folder of images for each row-column pair. Filenames for cropped images automatically embed the row and column metadata provided (e.g., PlateName_RowLabel_ColLabel.png), with the resulting images packaged into a zip archive for download and downstream analysis.

These segmented images of individual observations can be loaded into the BLISS scoring app for blinded score assignment. The BLISS scoring tool hides all image identifiers, randomizes spot order via a Fisher-Yates shuffling algorithm, and presents spots for spot images for scoring on a 0-4 (default) or user-defined (optional) scoring scale. To quantify intra-rater reliability, each spot can be presented multiple times via a replication factor (a replication factor of two was used in case studies included here) that determines the number of times an image is to be scored. Scores are assigned by clicking on individual images to assign them the currently selected score. Images, once scored, are moved to the bottom of the list in score-order for faster assignment and the opportunity for blinded score correction for incorrect assignments. Agreement is summarized with Cohen's weighted $\kappa$ using quadratic weights. BLISS exports analysis-ready CSV files with one row per observation (spot) containing: unique spot identifier, score, group covariates, numeric predictor value derived from row/column data. The resulting exported file can be loaded directly into the analysis tools MICalculator and ordinalMIC described in Supplemental Methods S10 below. Plate-reconstruction with overlaid score assignments as well as scoring agreement metrics are also provided in the download zip file provided.

### S10 MICalculator and ordinalMIC

The ordinalMIC R package wraps the ordinal R package to compute MIC*, ∆MIC*, and ∆log2MIC* with delta-method confidence intervals and Wald p-values. Given a fitted clm model object and the name of the numeric covariate of interest (e.g., concentration) with optional transformation, it extracts $\hat{\boldsymbol{\theta}}$ to estimate MIC* estimates per group using equations in S3 and S5 above. Gradients and Wald statistics are generated from the model's covariance for marginal estimates. For models with interactions involving $x_c$, the general linear form is used. For non-symmetric link functions, the constant $C_{link}$ and any fitted scale effects are automatically incorporated into MIC* estimation.

The MICalculator web application utilizes serverless WASM and JavaScript functionality to provide identical functionality in a no-code environment, requiring only an internet connection. Uploaded CSV files must include one row per spot with columns for score, the numeric predictor on the experimental scale, and group covariates. The app fits the PO model via an ordinalMIC API, reports MIC*, contrasts, and confidence intervals. An Rmarkdown document is produced to maximize reproducibility. By default, scoring replicates are processed by random sampling per spot image prior to analysis.

### S11 Reproducibility Information

All code used for simulations and data processing is archived at Zenodo DOI provided in methods of the main text. The code used for MIC* analysis is provided in the R package ordinalMIC version 0.1.2. We used R version 4.4.1 and the following R packages: arrow v. 21.0.0, curl v. 6.2.2, doParallel v. 1.0.17, doRNG v. 1.8.6.2, emojifont v. 0.5.5, foreach v. 1.5.2, ggpattern v. 1.1.4, ggsci v. 3.2.0, gt v. 1.0.0, gtsummary v. 2.2.0, here v. 1.0.1, iterators v. 1.0.14, ordinal v. 2023.12.4.1, ordinalMIC v. 0.1.1, patchwork v. 1.3.0, rngtools v. 1.5.2, survival v. 3.8.3, tidyverse v. 2.0.0, VGAM v. 1.1.13.

### S12 Missing data and Random Effects

If scores are missing completely at random, PO fitting remains unbiased for regression parameters and thus for MIC*-based derivations. If missingness depends on observed covariates, rather than assigned experimental covariates, inclusion of those parameters in the PO model leads to reduced model bias. Clustered designs (e.g., biological replicates on different days) can be accounted for by including these variables as random effects (intercept, slope, or both as needed) in the fit PO model. The MIC* estimator and delta-method derived variance carry over using the mixed-model covariance.

### S13 MIC* framework flexibility

The MIC* framework is a special case of a more general family of model-derived endpoints that map ordinal responses to the experimental scale of a continuous predictor. In the main text, we defined MIC* as the concentration at which the fitted PO model predicts 50% probability of "no growth" (score $Y = 0$). These specific choices, score of 0 and probability of 0.5, work well for MIC calculation of microbial inhibition assays, but the same modeling approach can yield biologically meaningful endpoints at any ordinal category ($EC_{50}$) or probability level ($LD_{99}$).

Formally, for an $Y$ score threshold $k \in \{0,\ldots,J-2\}$ and any probability level $p \in (0,1)$, let $T^*_{k,p}(x_{nc})$ be the concentration $c$ such that $P(Y \le k \mid c, x_{nc}) = p$ . Under a cumulative-link model $g\{P(Y \le j)\} = (\alpha_j - \eta)/s$ with link $g(\cdot)$, cutpoints $\alpha_j$, optional scale $s > 0$, and linear predictor $\eta$, the solution on the modeling scale $x_c$ is

$$x^*_c(k,p;x_{nc}) = \frac{\alpha_k - x^*_{nc} - sC_{link}(p)}{\beta_c}, \quad C_{link}(p) \equiv g(p),$$

And the experimental scale endpoint is $T^*_{k,p} = f^{-1}\{x^*_c(k,p;x_{nc})\}$ with $f(c) = log(1+c)$ utilized in our analyses. MIC* corresponds to $T^*_{k=0,p=0.5}$ under a symmetric link function (so $C_{link}(0.5) = 0$). Absolute and relative contrasts, uncertainty via the multivariate delta method, and hypothesis testing follow exactly as in the MIC* case after replacing $\alpha_0$ with $\alpha_k$ and $C_{link}(0.5)$ with $C_{link}(p)$. Because the PO model is invariant to monotone relabeling and collapsing adjacent categories, $T^*_{k,p}$ is well defined so long as k refers to a valid cumulative boundary in the fitted model. Furthermore, PO models (with a symmetric link function) exhibit palindromic invariance, such that reversing the order of the scores only changes the signs of beta coefficients but not the parameter estimates.

Choice of $(k,p)$ should be driven by biology of the phenotype being scored and scientific questions. When scores encode increasing viability (e.g., 0 = no growth, 4 = maximal growth), k=0 targets the onset of any detectable growth ("inhibition, on average"), while k=3 targets a more stringent definition (e.g., "minimum required to observe any defect"). Probability levels near p=0.5 yield median-like endpoints that are stable and interpretable. More conservative endpoint choices can be made, like 0.9 if the goal is to location concentrations where zero scores are almost certain. Extremely high or low choices of p increase the risk of extrapolation beyond the observed range. Because of this, we utilize p=0.5 to obtain a median value that matches observed values. When domain standards exist, such as clinical category boundaries or biofilm grade cutoffs, aligning score cutoff k to the relevant boundary yields endpoints directly comparable to those standards.
The same flexibility applies to the predictor and link. Any monotone transformation $f(\cdot)$ of the monotonic predictor (time, temperature, multiplicity of infection, nutrient level), can be used for modeling so long as reasonable assumptions are met. The endpoint can always be returned on the experimental scale via $f^{-1}(\cdot)$.

This generalization enables endpoint choice tailored to diverse microbiological applications. For biofilm assays scored 0-3 across an antimicrobial gradient, $T^*_{2,0.5}$ can be interpreted as a median $EC_{grade\ge 2}$. For plaque assays with ordinal clarity ratings across MOI, $T^*_{1,0.8}$ identifies the exposure at which minimal plaque formation is very likely. In each case, absolute and relative contrasts of $T^*_{k,p}$ between groups quantifies treatment effects on the experimental scale with corresponding uncertainty estimates.

### S14 Study design

Because MIC* is identified by the point at which the linear predictor for the first cumulative boundary crosses the 50th quantile, observations near the transition region are most informative for estimating MIC*. The same sample size but with concentrations at linearly or logarithmically equally spaced scale increases the variance of the MIC* and ∆MIC* estimators. When prior knowledge is limited, serial dilutions of concentration can provide improved point estimates over classical observed MIC estimation approaches.

Effect sizes of biological interest can inform the degree of granularity warranted for testing.

### S15 Numeric stability and MIC* edge cases

The MIC* estimator assumes a monotonic relationship between the cumulative category probabilities and exposure over the reported range of concentrations. Put simply, the effect of concentration should be the same biological effect or phenomenon at high scores as low scores. Nonmonotonic responses (e.g., Eagle effect) can be addressed either by nonlinear modeling of $x$ or by defining MIC* with respect to a restricted exposure interval and explicitly reporting that interval. MIC* estimation is most defensible within the support of the observed exposures. When estimates lie beyond the highest tested concentration, they should be noted as such and, if appropriate, followed by experiments that extend the exposure series. A second possible error is failure of the PO model to fit. This most often occurs when a single group exhibits no score variability throughout the entire range of tested concentrations, which results in no ability for the model to estimate $\beta_c$.

### S16 Small-sample and resampling-based inference

Delta-method based confidence intervals are fast and accurate in moderate samples. In small samples, or where the concentration effect of the testing compound is weak, profile-likelihood or bootstrap procedures can improve CI coverage. Profile-likelihood CIs for MIC* can be obtained by inverting a likelihood-ratio test on the implicit equation for $x_c^*$. Namely, solve for $P(Y \leq 0) = 0.5$ at fixed $x_c^*$ while re-estimating remaining parameters. Parametric bootstrap explains asymptotic normality of $\hat{\boldsymbol{\theta}}$ to form confidence intervals for MIC* and tested contrasts. This non-parametric bootstrap approach should resample at the unit of randomization (plate or biological replicate) rather than individual spots, to maintain the design's dependency structure. Each bootstrap dataset is refit and contrasted to capture both model and sampling variation.

### S17 Reporting recommendations for uncertainty and multiple testing

Many anticipated applications of MIC* analysis involve the simultaneous testing of MIC* across multiple contrasts of strain, conditions, or compounds. Because all MIC* contrasts are smooth functions of the coefficients estimates from a single fitted PO model, their join large-sample distribution is approximately multivariate normal by the delta method.

Let $g_k(\theta)$ denote the $k^{th}$ MIC* contrast (ΔMIC* or Δ$\log_2$MIC*), where $\hat{\boldsymbol{\theta}}$ is the MLE of the PO model parameters and $\sum = Cov(\boldsymbol{\theta})$. Let $J$ be the Jacobian with rows $\nabla g_k(\hat{\theta})$, then

$$\hat{g} \;\approx\; N(g, J\Sigma J^{\top})$$

Confidence intervals for all contrasts can then be constructed simultaneously while accounting for coefficient correlations. In practice, three options for multiple test correction can be used. First, familywise error rate (FWER) can be controlled by applying Holm's step-down procedure to the individual Wald p-values computed from the marginal delta-method variances which is uniformly more powerful than the Bonferroni correction while maintaining equivalent error control. No additional computation is required for this approach, and it is robust to contrast dependence. Second, simultaneous inference can be based on the join MVN approximate by simulating draws from $N_{\mathrm{k}}(0, J\Sigma J^{\top})$ to estimate the distribution of $max\ |Z_k|$, where $Z_k$ are standardized contrasts. The empirical $(1-\alpha)$ quantile yields a single critical value that produces simultaneous confidence bands for all contrasts. Third, when K is large and discoveries are expected to be sparse, error control using false discovery rate (FDR) via Benjamini-Hochberg applied to individual Wald p-values is appropriate for screening while acknowledging a small expected proportion of false positives. For complex dependence structures or small-sample settings, a Westfall-Young permutation procedure can be implemented by permuting group labels within blocking factors, refitting the PO model, recomputing the vector of MIC* contrasts, and using the empirical max-type null distribution to adjust p-values. This approach also preserves correlation patterns induced in the design and is more computationally intensive.
